# Drug Response Prediction and Biomarker Discovery Using Multi-Modal Deep Learning

**DOI:** 10.1101/2023.11.16.567479

**Authors:** Farzan Taj, Lincoln D. Stein

## Abstract

A major challenge in cancer care is that patients with similar demographics, tumor types, and medical histories can respond quite differently to the same drug regimens. This difference is largely explained by genetic and other molecular variabilities among the patients and their cancers. Efforts in the pharmacogenomics field are underway to understand better the relationship between the genome of the patient’s healthy and tumor cells and their response to therapy. To advance this goal, research groups and consortia have undertaken large-scale systematic screening of panels of drugs across multiple cancer cell lines that have been molecularly profiled by genomics, proteomics, and similar techniques. These large data drug screening sets have been applied to the problem of drug response prediction (DRP), the challenge of predicting the response of a previously untested drug/cell-line combination. Although deep learning algorithms outperform traditional methods, there are still many challenges in DRP that ultimately result in these models’ low generalizability and hampers their clinical application. In this paper, we describe a novel algorithm that addresses the major shortcomings of current DRP methods by combining multiple cell line characterization data, addressing drug response data skewness, and improving chemical compound representation. The result is an open-source, Python-based, command-line program available at https://github.com/LincolnSteinLab/MMDRP.

## Introduction

Cancer subtypes differ at the pathway activity level, their patterns of clinical progression, and their response to radiation, immunotherapy, and chemotherapy^1^. However, due to inter-patient molecular heterogeneity, even patients with the same cancer subtype have widely varying responses to the same therapy. This heterogeneity is thought to be primarily due to a combination of germline genetic differences among patients and somatic mutational variations within their tumors^2^.

Over the past decade, substantial technological advancements in biological profiling have given rise to the high throughput ‘omics’ era of biology, revolutionizing our understanding of cancer and many other diseases. Going beyond research insights, ‘omics technologies have opened the door to identifying molecular biomarkers that predict when cancer will respond to a particular therapy or when a patient is at risk of having an adverse reaction to a therapy^3^. This is the goal of precision oncology, which seeks to match a patient and their tumor to the therapy most likely to benefit them^4^. Pharmacogenomic screening of cancer cell lines has emerged as a critical approach for understanding the role of biological background in the sensitivity to therapeutics. Like other pre-clinical models, cell lines do not perfectly simulate cancer in patients. However, by systematically measuring the change induced by different drugs on the viability or growth rate of a large panel of cell lines, these studies allow us to understand better how variations in the genome alter treatment response. In addition to discovering predictive biomarkers that help better match drugs to patients, these efforts can help guide the development of new drugs.

Unfortunately, the cross-product of all promising compounds, cell lines, and genetic backgrounds is too large to evaluate empirically. For this reason, there is considerable interest in using the data from high-throughput cell line screening studies to build computational models capable of predicting drug efficacy in untested cell lines and compounds. This task is referred to as drug response prediction (DRP), and it is hoped that the ability to predict cell line response to drugs will be a stepping stone to predicting drug response in patients. In 2014, the USA’s National Cancer Institute (NCI) sponsored a DREAM community competition for DRP^5^. One key insight emerging from this competition was that those models that learn complex and non-linear (e.g., non-additive) patterns within the data outperform those that do not. Another is that some models leverage one data type better than others. Thus, applying the same algorithm to multiple data types may not be ideal. A final insight from this challenge is that different representations of the same data type can provide improved predictive capabilities. Despite the prominence of deep learning in many fields, none of the approaches in NCI-DREAM’s challenge applied neural networks. Since then, new programmable deep learning frameworks such as TensorFlow^6^ and PyTorch^7^, new cell line characterization datasets such as the Cancer Cell Line Encyclopedia^8^ (CCLE), and new drug response datasets such as the Cancer Therapeutics Response Portal version 2^9^ (CTRPv2) and Genomics of Drug Sensitivity in Cancer version 2^10–12^ (GDSC2) have been published^13–14^. In this report, we apply the insights from previously published studies to the new drug response data sets to develop a Multi-Modal Drug Response Predictor (MMDRP), a Python-based program that uses a multi-modal neural network to predict the efficacy of drugs on cell lines. Some of the innovations of this model include (1) the assignment of stronger weights to less frequently observed samples during training, which prevents overfitting to the prevalence of ineffective drug and cell line combinations; (2) a modular, multi-modal framework that can take advantage of multiple omic data types simultaneously for better predictions, without requiring that they all be present throughout the training data set; (3) a graph neural network for drug representation, which allows for a more informative representation of drug molecules and their structure; and (4) a final step in which the multi-modal input is combined using a suitable fusion method, which combines the complementary information from various omic sources. We assessed the model’s performance under various scenarios using a set of custom cross-validation schemes, which revealed the benefits of the proposed methods compared to previous approaches and illuminated the general strengths and weaknesses of the model. This work also assesses the utility of previously unused cell line profiling data for DRP. These results act as a guideline for future work in DRP.

## Methods and Results

### Training and testing datasets

To train and test MMDRP, we use used the following prominent pharmacogenomic datasets:

- Cancer Therapeutics Response Portal version 2^9^ (CTRPv2) – A dataset containing the sensitivities of numerous cancer cell lines to small-molecules.
- DepMap^31,32^ – An online database that hosts the Cancer Cell Line Encyclopedia, which contains various omic profiling data on more than a thousand cancer cell lines.

Exploratory data analysis of these datasets found challenges in data quantity, quality, and coverage. Two issues emerged. First, the molecular profiling data is sparse in the sense that different cell lines have been profiled for molecular characteristics that do not completely overlap (**Figure 1a**). Furthermore, the distribution of prediction targets, mainly the area above the dose-response curve (AAC), is non-uniform (**Figure 1b**). Although this reflects real-world pharmacology, the data skewness can hamper both the training and the testing of any model and should be given extra consideration when designing a model.

**Figure 1.**
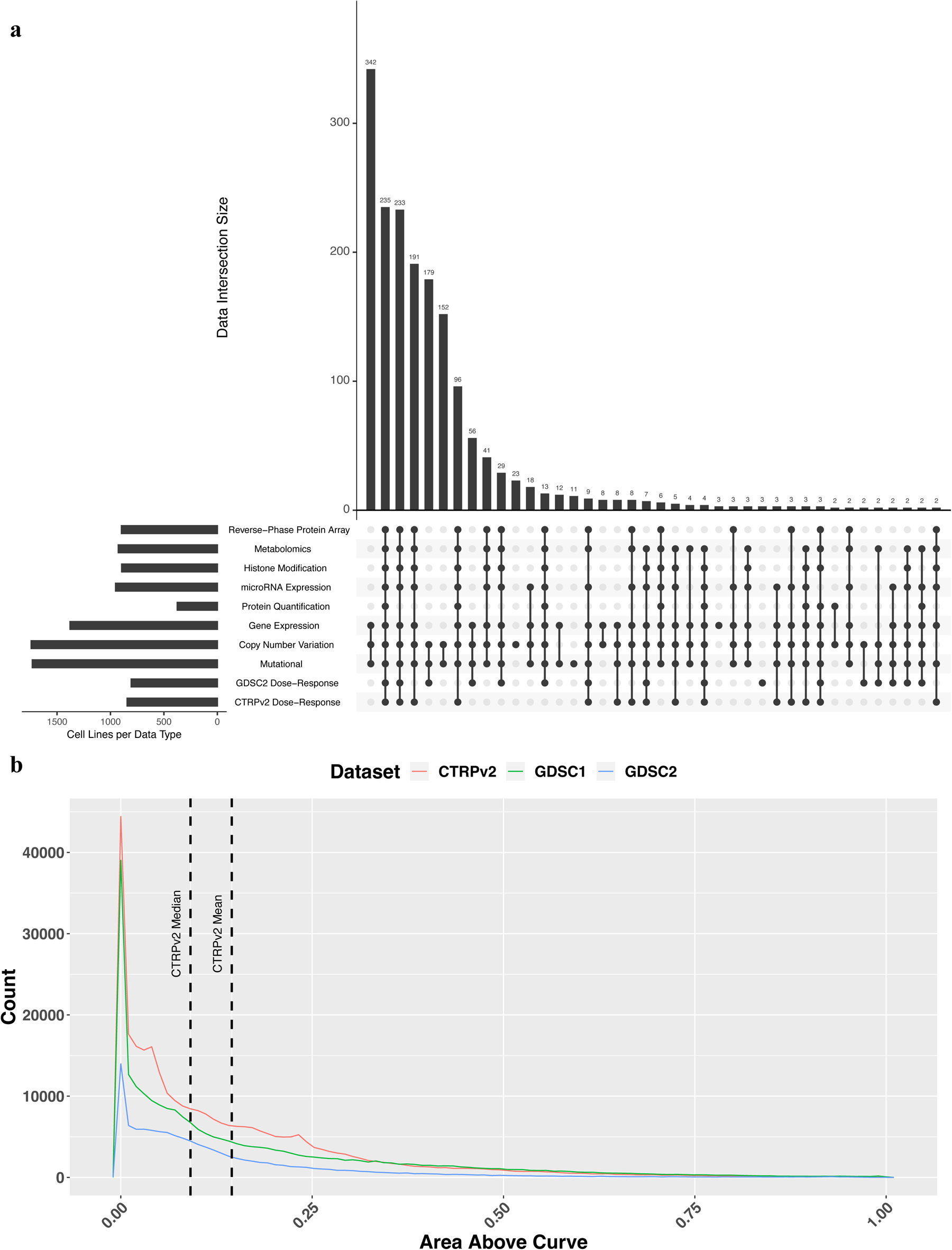
Exploratory data analysis of DepMap, CTRPv2, and GDSC2 datasets. (**a**) UpSet plot depicting the overlapping cell lines by data type from DepMap, CTRPv2, and GDSC2 datasets. Overlaps are ordered by the size of the intersections among sets. The top intersection of cell lines with gene expression, copy number variation, and mutational data with 342 cell lines does not overlap with dose-response data from the CTRPv2 or GDSC2 datasets. Number of cell lines in each dataset, from the top: RPPA (n=898), Metabolomics (n=927), Histone Modification (n=896), microRNA Expression (n=954), Protein Quantification (n=378), Gene Expression (n=1378), Copy Number (n=1742), Mutational (n=1731), GDSC2 (n=806), CTRPv2 (n=844). (**b**) Distribution of areas above the dose-response curve (AACs) from three drug response data sets. Higher AACs indicated a stronger response. In CTRPv2, AAC has a mean of 0.145 and a median of 0.091 (vertical dashed lines). As shown, most cell line-drug combinations in all three dose-response datasets have an AAC closer to zero. The higher ranges of AACs above 0.5 are less well represented. (CTRPv2: n=310,792, GDSC1: n=244,247, GDSC2: n=115,732)

The major challenges to effective DRP are data quantity, quality, skewness, and completeness. To overcome these challenges, we devised a modular, multimodal neural network that first independently learns from each type of cell line profiling data and then intelligently combines these learned representations to predict the responses of each cell line to each tested drug. This approach overcomes the central challenge of data sparsity. It allows the algorithm to learn across data sets in which cell line/drug combinations have different subsets of molecular profiling data.

### Pipeline Design

The pipeline is divided into three sections: preprocessing, training, and validation, where the goal is to predict the drug response target. We embed these steps in a hyperparameter optimization workflow, searching for more optimal design characteristics of the omic processing modules rather than choosing arbitrary hyperparameters (**Figure 2**). We use area above the dose response curve (AAC) as the prediction target because it has been shown to improve the predictive accuracy of tested models^16^, by capturing more information from the experiment than the more commonly used score, the IC_50,_ which is the half-maximal inhibitory concentration. IC_50_ has not been experimentally observed for many cell line and drug combinations within the tested dose range in drug response datasets. Furthermore, IC_50_ fails to differentiate two drugs with the same half-maximal inhibitory concentration but where one drug has a higher inhibitory power at a higher or lower dose^17^.

During preprocessing, data in each cross-validation training and validation split is standardized based only on the training split’s mean and standard deviation. Because of the data skewness problem, most available data are heavily skewed toward ineffective drugs with low AACs, and a minority have higher AACs. To mitigate this problem, we apply Label Distribution Smoothing^18^ (LDS) an algorithm that biases the selection of samples in the training set towards those that are less frequently seen by assigning a weight inversely proportional to the frequency of the drug response target to each sample in the training set, thereby forcing the model to learn across the entire range of values (**Supplementary** Figure 1).

In the training step, we train separate autoencoders to compress their respective molecular profiling inputs into space-efficient latent representations that capture the essential features needed to reconstruct the original data. In parallel, we apply a drug processing module to create a latent representation of each drug’s properties. In previous approaches, drug molecules were represented as Extended-Connectivity Fingerprints^15^ (ECFP), which are heavily engineered and precomputed features. This has the unwanted side effect of reducing the model’s ability for task-specific learning. To improve the representation and extraction of information from drug data in the context of DRP, we used a graph neural network (GNN) known as the AttentiveFP^19^ model. GNNs are a variant of neural network architectures that are well suited to model structured data such as networks and molecules. The AttentiveFP GNN models the physiochemical properties of molecules, which is an advantage over the traditionally used molecular fingerprints such as ECFP. Furthermore, the AttentiveFP model has learnable parameters that allow for context-specific learning. In contrast, molecular fingerprints do not allow for learning from physiochemical properties but rather describe the structural properties of the molecules they encode. Note that we use the published AttentiveFP model as is and have not tuned its hyperparameters to optimize it to the DRP task.

The latent representations of the cell line’s molecular profile must then be combined with each drug’s latent representation. Low-rank multimodal fusion^20^ (LMF) is a technique for combining multiple modalities in a neural network such that the latent representations of different features are forced to ‘interact’ with each other. LMF has shown higher performance than other fusion methods, such as (weighted) summation and simple concatenation. It is especially important for modeling biology since it is known that various biomolecules in the cell interact with each other and thus must also be allowed to interact when modeling biology in silico. We use LMF as the fusion method, and the output from this fusion is then passed to another final module, which predicts the AAC. We use the Root Mean Squared Error (RMSE) loss for model training.

## Evaluatation of Model Performance

To assess the model’s performance, we applied multiple fivefold cross-validation schemes. In each scheme, data were split in a manner to prevent the reappearance of specific samples from the training set in the validation set. The four splitting strategies were: 1) Split by Cell Line, 2) Split by Drug Scaffold, 3) Split by Both Cell Line and Drug Scaffold (simultaneous), and 4) Split by Cancer Type. Each of the four splitting strategies was done in a way that ensures that the members of a splitting criterion (e.g., cell lines) are not shared among the training and validation sets. Each splitting strategy results in a separately trained model that is then evaluated independently. To avoid leakage between drugs with similar structures, we also split on the drug scaffold, which is the backbone of a drug molecule, crucial in determining a molecule’s biological activity. This is more stringent than splitting on the drug name. The validation samples from each of the 5-folds were then combined to report a final total validation performance.

**Figure 2.**
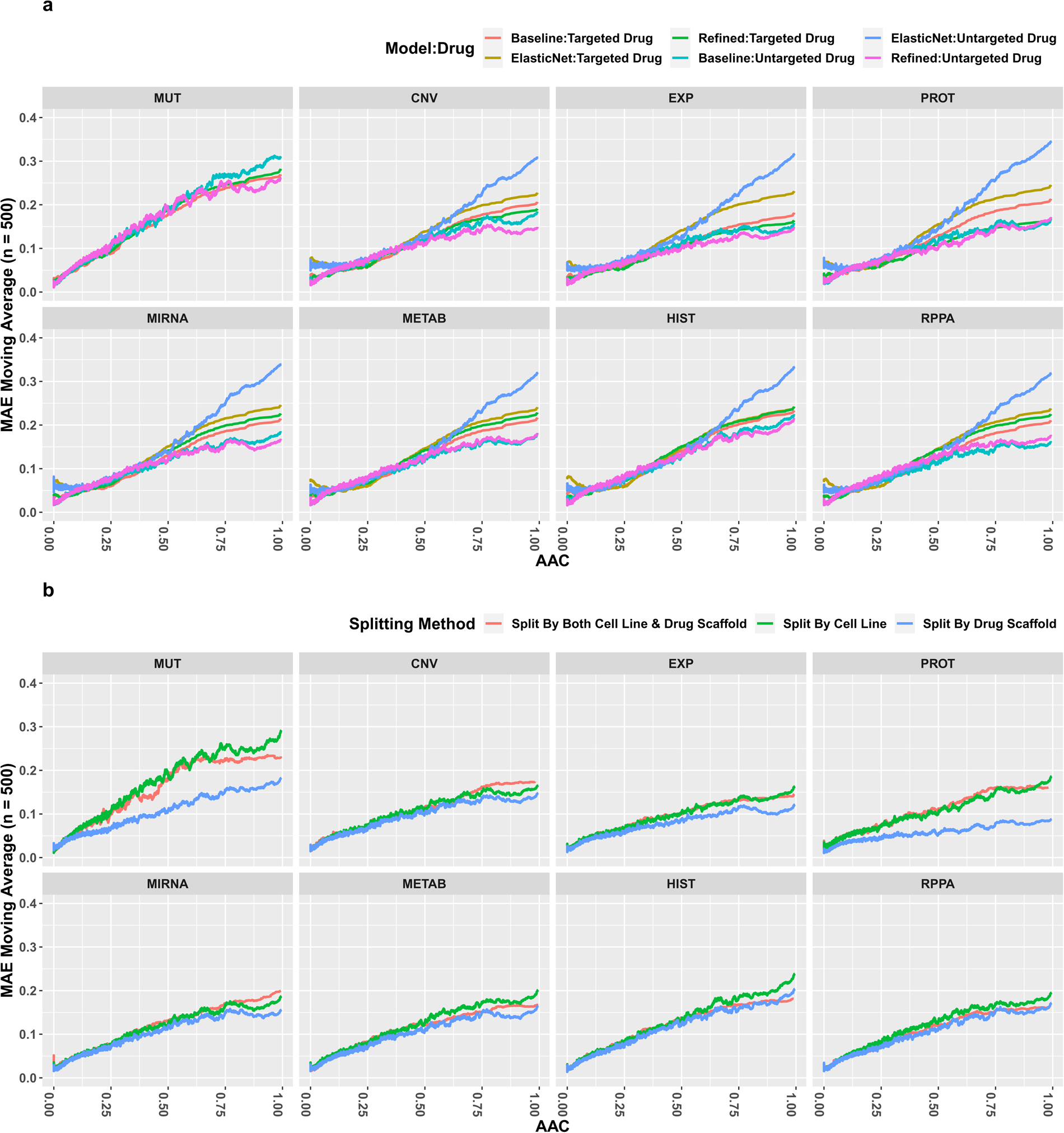
Hyperparameter optimization and training scheme for omic modules of multi-modal DRP models. We first run 40 hyperparameter search trials for each omic-specific autoencoder, using the aggregated cross-validation losses as a measure of optimality. After identifying an optimal hyperparameter configuration for each autoencoder, we then train each one on all available omics data. Then, we use the encoder submodel and connect it with the DRP submodel, and run 40 trials for hyperparameter search, in this case only searching for hyperparameters for the DRP submodel, but also for each cross-validation splitting scheme. Finally, we train and validate the model using different splitting schemes. Optimal hyperparameters are available on the GitHub repository.

Due to data sparsity, autoencoder sub-models using different molecular profiling data types each received different amounts of input data. To ensure fair comparisons, we create a restricted dataset that only contains data from cell lines with all eight molecular data profiles, i.e., the subset of complete samples. This subsetting reduced the initial dataset of 35 cancer types across 1776 cell lines to 24 cancer types across 331 cell lines (**Supplementary** Figure 2). This resulted in 125,526 data points, 40.4% of the original dataset of 310,792 data points.

Since drug response data is heavily skewed towards non-responsiveness, reporting a single performance score across the entire AAC range (e.g., root-mean-squared error or RMSE) is misleading. We used the trailing moving average of the mean absolute error (MAE) of 500 samples along the entire AAC range to report and compare model predictive accuracy. This assessment was absent from previous studies. In addition, we use RMSE for the performance measurement of a group of samples. Note that the RMSE of a single prediction is equivalent to its MAE and that lower scores are better for both MAE and RMSE.

### Bimodal Model Performance

We first evaluated the model performance when the drug structure and a single molecular profiling type (e.g. RNA expression) were provided. We refer to these as “bimodal models.” Most published methods in DRP use IC_50_ as the prediction target^13^, which prevents direct comparison of their results to those obtained with MMDRP. As the modification of previously published deep learning methods in DRP to use AAC instead of IC_50_ is non-trivial, we used a simple yet effective linear algorithm, the elastic net regularized linear regression algorithm, as our comparison control. We then compared the performance of elastic net against two implementations of MMDRP: 1) a baseline multi-modal neural network that does not employ the LDS, LMF, and GNN techniques described above referred to as MMDRP-base; and 2) a neural network using all three techniques, hereby referred to as the MMDRP-refined model. The changes in performance for each method are reported in the supplementary materials. **Figure 3a** compares the predictive performance of the baseline elastic net model, the baseline bimodal model, and the MMDRP-refined bimodal model. The MMDRP-refined bimodal models consistently outperform the baseline bimodal models in both targeted and untargeted drugs across the entire AAC range. The “split by drug scaffold” splitting method has lower losses than other splitting strategies (**Figure 3b**). Interestingly, the bimodal model with protein quantification data (PROT) performs significantly better in the “split by drug scaffold” scheme.

All models are generally better at predicting cell-line/drug combinations with lower AACs than higher AACs, a trend which remained consistent across the different models and splitting methods. We expect this because of the strong skew towards low AAC values across the training data.

Another observation is that samples treated with targeted drug therapies (defined in **Supplementary Table 1**) are harder to predict across the entire AAC range, shown in red and cyan in **Figure 3a**. Targeted drugs, on average, have higher AACs in the CTRPv2 dataset (**Supplementary Figure 3**). Furthermore, only 31 of the 481 drugs in the CTRPv2 dataset are untargeted, and such imbalance can affect the predictive performance of the models.

**Figure 3.**
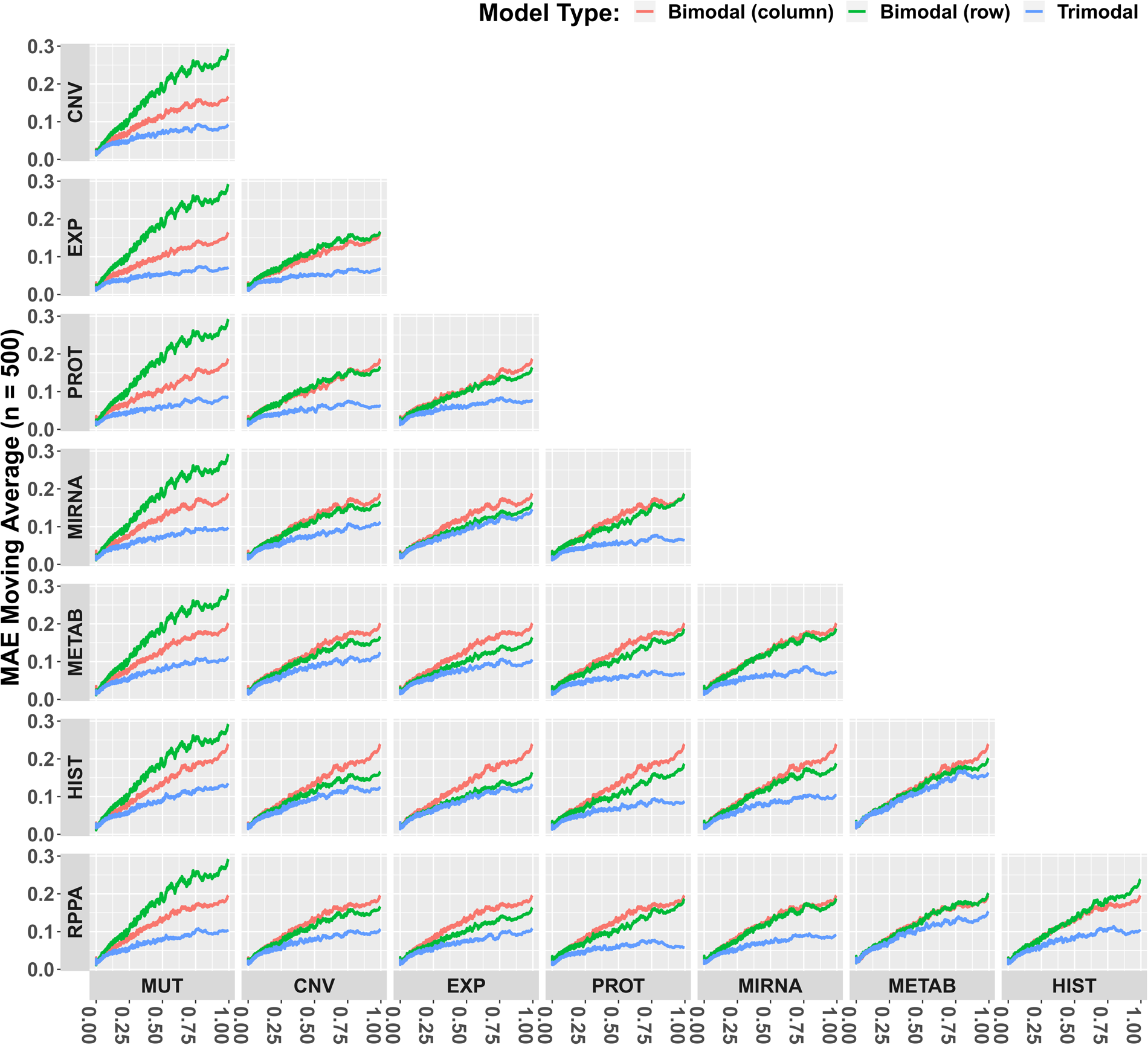
Predictive performance of bimodal models. (**a**) Comparison of the elastic net, baseline bimodal, and MMDRP-refined bimodal models. (MUT: Single Nucleotide Variation, CNV: Copy Number Variation, EXP: Gene Expression, PROT: MS-based Quantitative Proteomics, MIRNA: miRNA expression, METAB: metabolite abundance, HIST: Histone H3 modification, RPPA: Reverse-Phase Protein Array-based protein quantification) The use of all three methods (LDS, LMF, GNN) results in better performances, especially in the higher AAC ranges and in targeted drugs. Targeted drugs are harder to predict than untargeted drugs, especially at higher AACs. The model trained on gene expression data performs the best for samples with targeted drugs, followed by the models trained on RPPA, protein quantification, and metabolomics data. However, the model trained on RPPA data performs the best for untargeted drugs following the model trained on gene expression data. The mutational model has the lowest performance in both targeted and untargeted drugs. (**b**) Comparison of MMDRP-refined bimodal model performance across three splitting methods. (Lower MAEs are better)

Unexpectedly, the models’ drug response predictions based on cell line mutational data alone had the poorest performance of all the molecular data types. This was particularly evident in the bimodal model. Furthermore, there was less consensus on the predictions among the models in samples with higher AACs than those in the lower AAC range (**Supplementary Figure 4**). This shows that models trained with different molecular data types may be more suitable for particular cell line and drug combinations than others. The best models based on total RMSE losses by omic type, splitting method, and drug type are reported in . No single model consistently outperforms others in all scenarios.

### Multi-modal Model Performance

To assess the benefits of combining different molecular profiling data types for drug response prediction, we began by creating the trimodal case (drug structure plus two types of molecular profiles), resulting in 28 profile combinations for baseline and MMDRP-refined model configurations. As in the bimodal case, the trimodal MMDRP-refined models consistently outperform the baseline models in targeted and untargeted samples (**Supplementary Figure 5**). More importantly, MMDRP-refined trimodal models consistently outperform bimodal models across all data types (**Figure 4**). In addition, among the different splitting schemes, splitting by cancer type often has the highest losses in this subset, meaning that the generalization to novel cancer types is the most challenging task among the four splitting methods (**Supplementary Figure 6**).

**Figure 4.**
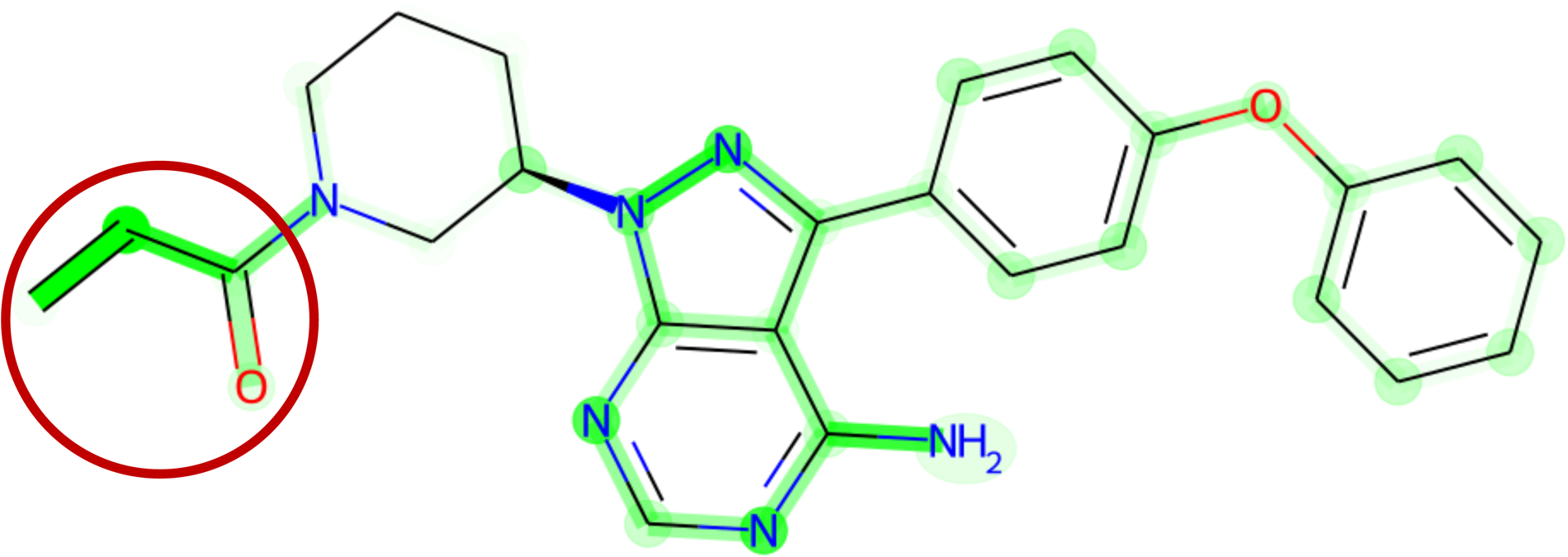
Trimodal vs Bimodal model performances across the AAC range. In all cases, the model that uses a combination of the two omic data types from the row and columns performs better than the model that uses an omic data type individually. (Lower MAEs are better)

Since the total number of possible molecular profiling data combinations increases for higher-order combinations, we selected data types from top-performing trimodal models and several complementary data combinations for testing. summarizes the total RMSE losses of the models tested. As in the trimodal case, we found that the majority of the highly performing models used the MMDRP-refined configuration.

Comparisons between MMDRP-refined quadmodal models to their trimodal and bimodal counterparts in the split by cell line scheme reveal that the trimodal models consistently outperform the three quadmodal models (**Supplementary Figure 7**). Within the current configurations tested, trimodal models often outperform higher order combinations. To assess the relevance of profiling data types for each drug-cell line combination, we measured the frequency of profiling data types used by models that predict AAC most accurately for each data point (**Supplementary Figure 8**). Note that the frequencies of the best model for each data point do not always correspond with the model that has the lowest total RMSE loss.

In summary, these results have shown that the combination of multiple complementary profiling data types can improve drug response predictions, whereas the combination of non-complementary data types can hurt predictive performance. Furthermore, improving data processing and learning by employing techniques such as the LDS, LMF, and GNN can dramatically improve learning from the incomplete and skewed drug response data, which can then improve the quality of the predictions in more clinically relevant cell line/drug combinations with higher AACs, as well as targeted therapies.

## Predicting Activities of Drugs with Repurposing Potential

Not infrequently, a drug that has previously been brought to market for the treatment of one disease is found to have utility in the treatment of a different disease. To measure MMDRP’s ability to identify drugs with repurposing potential, we focused on FDA-approved drugs within the CTRPv2 dataset. First, we found the subset of drugs that have higher true AACs in cancer types other than those for which they were approved. Then, we asked whether each model, having seen that cell line for the first time, could predict this higher AAC. **Supplementary Table 4** is a table of 76 potentially repurposable drug-cell line combinations that 1) had an AAC at least 0.2 higher in a non-cognate cell line as compared to all the cell lines derived from the tumor type for which the drug is typically prescribed, and 2) a bimodal MMDRP-refined model accurately (MAE ≤ 0.2) predicted this higher AAC, having seen the cell line for the first time (for the complete list, please refer to the GitHub repository). Interestingly, out of the 76 potentially repurposable drugs-cell line combinations, 48 of them were predicted with a model that did not use gene expression data. Here, we will focus on a typical example of a potentially repurposable drug.

Ibrutinib is a tyrosine kinase inhibitor, originally designed to target Bruton’s tyrosine kinase (BTK). It is FDA-approved for Chronic Lymphocytic Leukemia (CLL) and a few subtypes of lymphoma. In the CTRPv2 dataset, ibrutinib’s highest AAC in tested leukemia cell lines is 0.496. Interestingly, there are five breast cancer cell lines with similar or higher empiric AACs in CTRPv2. Trimodal MMDRP-refined models trained on data split by cell lines accurately predicted the sensitivity of all five cell lines to ibrutinib (). Additional validation of these predictions comes from recent research which confirms both the empiric and predictive results that ibrutinib treatment can inhibit breast cancer progression and metastasis in in-vivo mouse models^21^.

Although MMDRP has good predictive power, its potential utility as a diagnostic test in its current format is low due to the multiple molecular assays that would need to be performed on patient tissues or derived cell lines. Therefore, we assessed the model’s ability to identify the most informative features driving a prediction for biological interpretation and biomarker discovery. The Integrated Gradients method helps identify features that are most salient for a model’s predictions. It works by integrating the gradients of the model’s output with respect to the input along a straight path from a baseline input (usually all zeros) to the actual input. This gives a detailed decomposition of how much each feature contributes to the prediction; these contributions are referred to as attributions. For biomarker selection, we focused on attributions for omic features (e.g., CNV/gene expression features). Since the attributions are unique to each input sample, we focused on samples with higher AAC (more efficacious drugs). Although we focused on omic features, we can also make attributions to drug features.

We used the Integrated Gradients method to interpret the CNV + EXP MMDRP-refined model for Ibrutinib+EFM192A. This model predicts an AAC of 0.51 for Ibrutinib in the EFM192A cell line, close to the empiric AAC of 0.658. Nine of the ten features attributed with the highest influence on the model were all from gene expression data. Surprisingly, although Ibrutinib targets BTK, the model did not consider the expression of this gene important and gave low attribution scores in both gene expression and CNV data. BTK’s gene expression is in the 46^th^ percentile compared to all cell lines and the 48^th^ percentile in breast cancer cell lines, while its copy number is in the 93^rd^ percentile compared to all cell lines and the 81^st^ percentile in breast cancer cell lines (**Figure 5**). In contrast, of the nine genes selected by the algorithm, DERL1, and FAM91A1’s expression levels are in the 99^th^ percentile in the EFM192A cell line compared to other breast cancer cell lines and display correlation with breast cancers’ response to ibrutinib, including EFM192A (**Figure 5**). Recent studies have identified Derlin-1 (the protein product of DERL1) as a growth promoter in breast cancer, and patients with a high expression of Derlin-1 were found to have a significantly lower prognosis than patients with a low expression of Derlin-1^22,23^. *FAM91A1* was recently identified as a candidate target gene for breast cancer risk signal^24^. In addition, interpreting the model’s GNN drug module using the same Integrated Gradients method reveals that the model correctly focuses on ibrutinib’s acryloyl group when predicting its effectiveness on the EFM192A cell line (**Figure** 6). Together, these results suggest that the algorithm is capturing biologically and chemically relevant response features.

**Figure 5.**
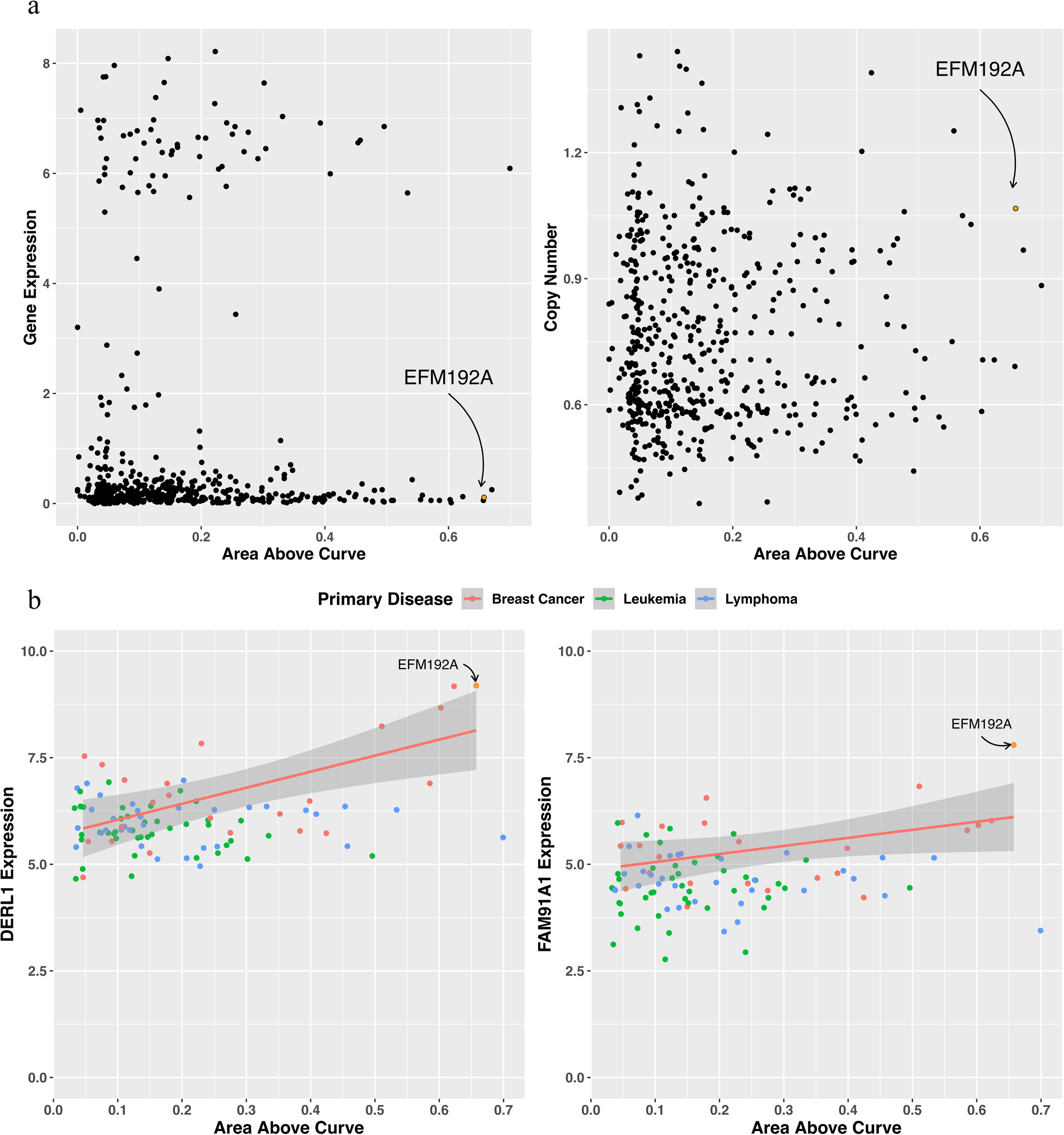
Cell Line Response to Ibrutinib. **(A)** Empiric response to Ibrutinib in cell lines from CTRPv2. (Left Panel) Relationship between BTK gene expression level (y-axis) and drug response AAC (x-axis). (Right Panel) Relationship between BTK gene copy number (y-axis) and drug response AAC (x-axis). The highly sensitive breast cancer cell line EFM192A is highlighted. **(B)** DERL1 and FAM91A1 relationship with drug response in ibrutinib-prescribed cancer cell lines. Among all breast cancer, leukemia, and lymphoma cell lines, DERL1 and FAM91A1’s expression show higher correlation with AAC in ibrutinib-prescribed cancer cell lines than other cell lines (Adjusted R-squared 0.3505 and 0.1336 respectively).

**Figure 6.**
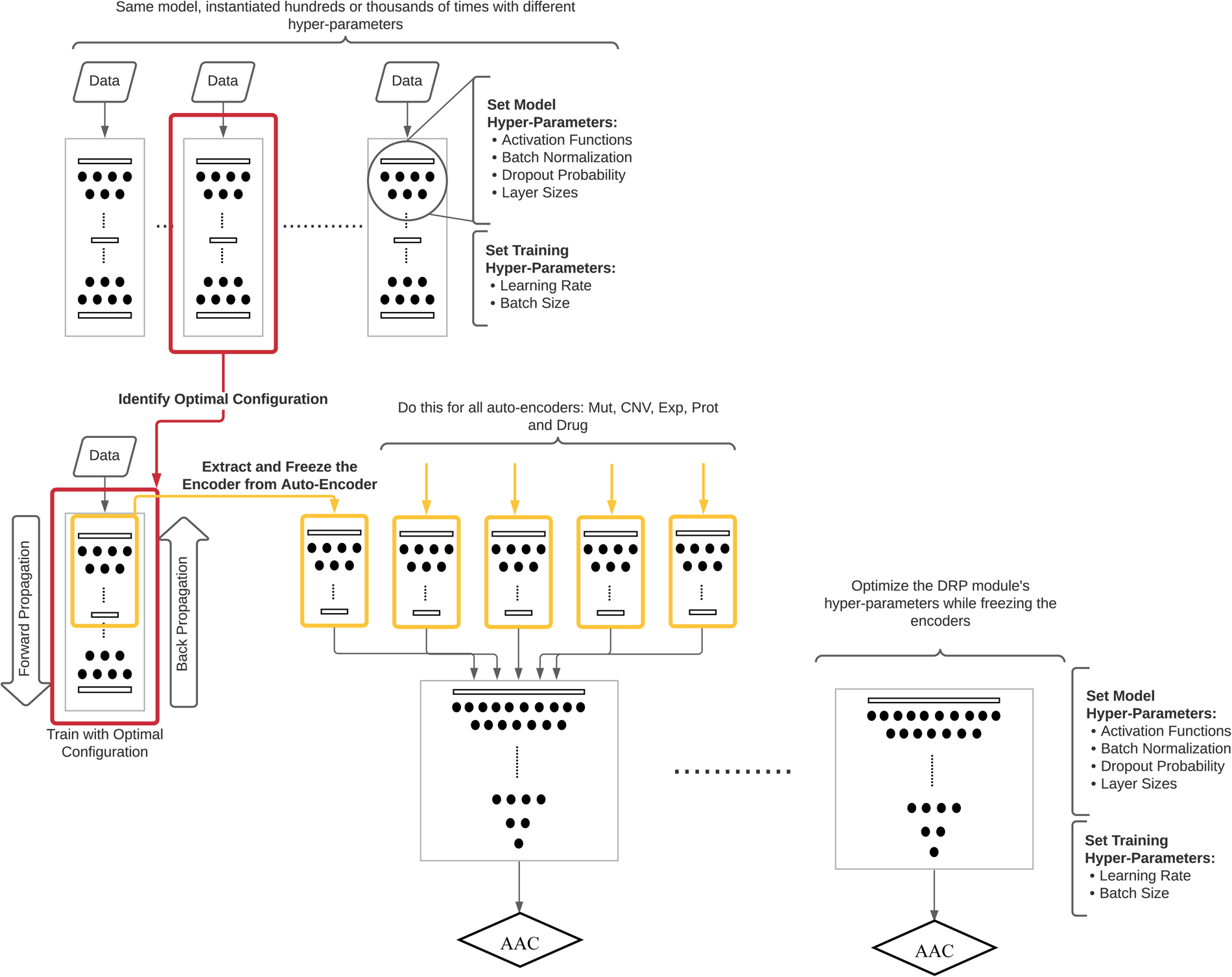
GNN’s attributions on Ibrutinib used on the EFM192A1 cell line. The densely highlighted regions correspond to higher attributions, whereas lighter colors correspond with lower attributions. The carbon in the circled acryloyl functional group, highlighted with a red circle, forms a covalent bond with the C481 residue on the BTK protein. The model correctly identifies this functional group as important. For better visualization, refer to the PDB entry 5P9J. The same functional group may bind to similar active sites in other cancer-related kinases.

## Discussion

Here, we describe MMDRP, a multi-modal drug response prediction model that uses multiple varieties of cell line molecular profiling data and a graph-based representation of drugs to predict the activities of drugs in cancer cell lines. As measured by MAE and RMSE on drug/cell line combinations that present in the training set, MMDRP is markedly more accurate than a baseline predictive model constructed using the elastic net framework. A direct comparison with other deep learning-based predictive models was not possible, as they are trained to predict IC_50_ rather than MAE. Our experiments have rigorously evaluated the applicability of multiple ‘omic data types, both singly and in combination, for drug response prediction for the first time.

We readily acknowledge the limitation of cell line studies to predict the efficacy of a drug in human patients. Indeed, the same caveats apply to other pre-clinical models, such as organoids and mouse models. Many drugs that seemed promising at the pre-clinical phase fail during clinical trials. However, it is important to remember that a drug cannot advance to a clinical trial unless it has shown promise in pre-clinical studies, and pre-clinical models represent a bottleneck in the drug discovery pipeline. If we can accurately predict which drugs are most likely to be effective in pre-clinical models, then we can reduce this bottleneck by increasing the pre-clinical pass rate. Hence, drug response prediction algorithms are useful for drug discovery even if they do not directly advance the precision oncology goal of predicting the response of individual patients to therapy.

Our work has identified several issues in the datasets available to DRP researchers. One problem is the difficulty in making useful predictions from clinically relevant cell line and drug pairs that are underrepresented in the training set. This is particularly acute for molecularly targeted drugs, which tend to have fewer drug/cell line pairs in drug response screening datasets than their older cytotoxic counterparts. Results have indicated it is still possible to improve predictive performance using existing cell line data for DRP through better modelling approaches. The use of label distribution smoothing (LDS), low-rank multimodal fusion (LMF), and a graph neural network (GNN) for drug representation significantly improved the performance across the AAC range. Lastly, interpreting performant models allows for biomarker discovery and drug repurposing.

It has been common for previous DRP approaches to solely use gene expression data. This work has shown that other cell line profiling data types are not only useful for making drug response predictions but also for identifying drugs with repurposing potential and the discovery of novel biomarkers. Furthermore, combining two or more profiling data types significantly improved performance over individual use of data types, especially when the LMF fusion method was used. Based on results from the trimodal models, CNV and gene expression data seem to capture cell line biological knowledge better and allow the model to generalize better to novel cell lines, whereas protein quantification data (PROT & RPPA) better capture cell line and drug interactions such that the model can generalize better to novel small molecules, especially targeted therapeutics. Mutation data was generally ignored by the best-performing classifiers, perhaps due to its representation as binary data.

Unexpectedly, higher-order combinations of molecular profiling data types had poorer performance than the trimodal models. It is unclear whether the added information provided redundant information or, alternatively, whether the model could not incorporate the new knowledge to make accurate predictions. This depends on the different cancer-related pathways that are dysregulated in different cancers, which may be better reflected in a specific group of macromolecules than others. We previously noted that the frequencies of the best model for each data point do not correspond with the model with the lowest total RMSE loss. For example, although the model that uses protein quantification data has the lowest total RMSE in the ‘split by drug scaffold,’ a larger proportion of data points are predicted more accurately by models that use gene expression data. Since no single model has the best predictive performance on all samples simultaneously, we recommend predicting the AAC of a new sample using all reported models and only consider further laboratory assessment if at least a few models predict an AAC in the useful range. Fortunately, performing predictions using neural network models is much faster than training them.

The four cross-validation schemes we used in this project gauge the models’ performance in typical industrial and clinical settings. For example, the objective may be to repurpose an approved drug for novel cancer types or subtypes where the evaluation by splitting by cancer type or cell line would be more appropriate. In contrast, when designing a novel drug, splitting by drug scaffold would be appropriate. As discussed before, most previous work in DRP has failed to assess generalization performance to novel cell lines, drugs, and cancer types. As molecular testing technologies advance, the difference among patients with cancer of the same body sites becomes clearer. Therefore, generalization to novel cell lines from familiar cancer types is perhaps the most relevant task to precision medicine. However, results also show that generalization to novel drug scaffolds and targeted drugs is also difficult. The latter results are in contrast with previous work published by Jang et al.^17^ in 2014, where it was concluded that models with pathway-targeted compounds are most likely to yield the most accurate predictors compared with those trained on broadly cytotoxic compounds. Therefore, future efforts in DRP depend on the expansion of current pharmacogenomic datasets such that they would encompass more cell lines and drug classes. Model-level engineering such as the incorporation of drug targets, protein structures, and pathway information can also help to enhance existing data.

In addition to cell lines, drugs, and cancer types, model generalization across the entire AAC, especially at higher AACs, is also vital. We are more interested in effective drugs than ineffective ones, although accurate prediction of the latter also has clinical utility. This task is hampered by the lack of sufficient data in higher AAC ranges and will require expanding existing drug sensitivity datasets.

Recent work on deep learning model interpretation allows us to understand the basis for neural network predictions. In this report, we applied the Integrated Gradients method to dissect model predictions for potentially repurposable targeted drugs and identified a series of predictive biomarkers that seem to be biologically relevant. In addition to the ibrutinib and BTK examples we report here, we have assembled a list of 76 predictions of effective cell line/drug combinations that can be used for drug repurposing and/or biomarker identification (**Supplementary Table 4**). In addition, complex modeling enables the discovery of composite biomarkers composed of the non-linear combination of multiple measurements^25^. Integrated Gradients assigns attributions to every single feature, and secondary analyses are required to identify the group membership of these features and their interaction to identify composite biomarkers.

MMDRP addresses some of the previous shortcomings of DRP, but certain challenges remain. Among several improvements that can be made to the algorithm, one would be to enhance the GNN by converting it to a graph-based autoencoder that could be pre-trained with the millions of available 2D compound structures currently available. In addition, different MMDRP models had the best predictive performance on different subsets of the data, and it is not clear which model’s predictions are the best for making drug response predictions on a new cell line or drug. To avoid testing multiple models, one approach would be creating a model that uses all the molecular data types while automatically subsetting input features and model parameters most relevant to its predictions based on the context of the given cell line and drug combination.

Recent work such as the Mixture-of-Experts^26^ (MoE) or attention mechanisms that allow neural networks to ignore inputs contextually^27^ can be used for this task. Furthermore, biomarker discovery from multimodal omic data can pave the way for multimodal, complex composite biomarkers with more predictive power than simple biomarkers. Although this work did not specifically assess multi-modal biomarkers, it serves as a necessary first step toward this new class of biomarkers. We believe these three improvements to the model will have the most immediate benefit both for increasing its predictive performance as well as its clinical utility.

Aside from algorithmic challenges, DRP data can also be improved in different ways. Cancer cell lines are not genetically stable and display clonal variation^28^. Single-cell profiling methods can improve pharmacogenomic screens by allowing us to associate drug sensitivities with each distinct subclone. It has already been shown that considering clonal heterogeneity can allow for a more accurate prognosis and drug sensitivity prediction^29^. In addition, we can identify minimal residual disease (MRD) cells that are not inhibited by the given drug to focus on drug development for those cells^30^. This data can also be used to retroactively assign cell type proportions to cell line profiling data, allowing us to add another layer of information for DRP. Single-cell profiling can, therefore, enhance DRP efforts to improve the discovery of personalized therapies.

In summary, drug response prediction is a clinically relevant problem that can be solved with current and emerging advancements in biological and computer sciences. The methods we have developed in this work have addressed some of the challenges and have allowed the identification of future work required to bring DRP closer to clinical use.

**Table 1.** Best multimodal models and their RMSE losses by omic type, splitting method, and drug type in the AAC ≥ 0.7 range. Highlighted cells correspond to the lowest RMSE for that column. (Lower RMSEs are better)

| Omic type (s) | Split By Both Cell Line & Drug Scaffold |  | Split By Cell Line |  | Split By Drug Scaffold |  |
| --- | --- | --- | --- | --- | --- | --- |
|  | Targeted Drug | Untargeted Drug | Targeted Drug | Untargeted Drug | Targeted Drug | Untargeted Drug |
| CNV + EXP + METAB | MMDRP-refined<br>0.312 | MMDRP-refined<br>0.189 | MMDRP-refined<br>0.257 | MMDRP-refined<br>0.159 | Baseline<br>0.559 | Baseline<br>0.235 |
| CNV + EXP + PROT | MMDRP-refined<br>0.395 | MMDRP-refined<br>0.215 | MMDRP-refined<br>0.36 | MMDRP-refined<br>0.222 | Baseline<br>0.578 | Baseline<br>0.268 |
| CNV + EXP + PROT + METAB | MMDRP-refined<br>0.45 | MMDRP-refined<br>0.197 | MMDRP-refined<br>0.364 | MMDRP-refined<br>0.174 | MMDRP-refined<br>0.528 | Baseline<br>0.233 |
| CNV + EXP + PROT + MIRNA + METAB + HIST + RPPA | MMDRP-refined<br>0.403 | MMDRP-refined<br>0.215 | MMDRP-refined<br>0.388 | MMDRP-refined<br>0.224 | MMDRP-refined<br>0.556 | Baseline<br>0.233 |
| EXP + RPPA + HIST + PROT | MMDRP-refined<br>0.4 | MMDRP-refined<br>0.218 | MMDRP-refined<br>0.358 | MMDRP-refined<br>0.223 | Baseline<br>0.594 | Baseline<br>0.245 |
| EXP + RPPA + PROT | MMDRP-refined<br>0.588 | MMDRP-refined<br>0.197 | MMDRP-refined<br>0.468 | MMDRP-refined<br>0.174 | Baseline<br>0.559 | Baseline<br>0.264 |
| MIRNA + METAB + HIST + RPPA | MMDRP-refined<br>0.471 | MMDRP-refined<br>0.196 | MMDRP-refined<br>0.41 | MMDRP-refined<br>0.173 | Baseline<br>0.546 | Baseline<br>0.228 |
|  | Targeted Drug | Untargeted Drug | Targeted Drug | Untargeted Drug | Targeted Drug | Untargeted Drug |
| MUT + CNV + EXP + PROT | MMDRP-refined<br>0.341 | MMDRP-refined<br>0.208 | MMDRP-refined<br>0.38 | MMDRP-refined<br>0.232 | Baseline<br>0.587 | Baseline<br>0.319 |
| MUT + CNV + EXP + PROT + MIRNA + METAB + HIST + RPPA | Baseline<br>0.516 | Baseline<br>0.389 | Baseline<br>0.635 | Baseline<br>0.352 | Baseline<br>0.578 | Baseline<br>0.288 |

**Table 2.** Ibrutinib in breast cancer cell lines and top models’ predictions (split by cell line).

| Cell Line | Lineage Subtype | Data Type(s) | True AAC | Prediction | MAE Loss |
| --- | --- | --- | --- | --- | --- |
| EFM192A | Breast Adenocarcinoma | METAB+RPPA | 0.658 | 0.595 | 0.063 |
| AU565 |  | EXP+MIRNA | 0.623 | 0.588 | 0.035 |
| SKBR3 | Breast Carcinoma | CNV+EXP | 0.603 | 0.586 | 0.017 |
| ZR7530 | Breast Ductal Carcinoma | HIST+RPPA | 0.585 | 0.585 | 0.000 |
| HCC1419 |  | MUT+RPPA | 0.511 | 0.502 | 0.008 |

## Implementation

### Software and Hardware

Neural networks were implemented using Pytorch ver. 1.7 and PyG ver. 1.7.2. Cross-validation schemes were implemented using sci-kit learn ver. 1.0.1. Model interpretation was performed using Pytorch Captum ver. 0.3.1. Hyperparameter optimization was performed using Ray [Tune] ver. 2.0.0.dev0. Model implementation as well as analysis scripts are available at: https://github.com/LincolnSteinLab/MMDRP.

### Datasets

#### Cell Line Characterization Data

This research utilized various cell line characterization data from the Cancer Cell Line Encyclopedia Project (CCLE) and DepMap^31,32^. In brief, this included point mutation and small insertion-deletion data, with an overall 1.27 million mutations identified across 1,750 cell lines. Specific gene mutations were selected for analysis based on their overlap with TCGA and COSMIC hotspots, resulting in a final tally of 8,772 genes represented as a binary vector. The work also involved using copy number aberration data and gene expression data, with gene-level copy numbers and gene-level expression values, respectively, utilized for this study. Protein quantification data, specifically high-throughput protein-quantification data from the Gygi lab, was utilized for some cell lines, focusing on approximately 5,000 proteins detected in all cell lines^33,34^.

Additionally, microRNA (miRNA) data, profiling the expression of 734 miRNAs across 953 cell lines, was also employed^32^. The metabolomics data and histone modification profiling data offered insights into changes in cell metabolism, and post-translational modifications of histone proteins were used to understand cancer mechanisms and cell characterization^32,35,36^ further.

Lastly, Reverse Phase Protein Array (RPPA) data was employed to measure protein expression levels for 214 proteins across 898 cell lines. See the appendix for more details.

#### Drug and Drug Response Data

In this study, drugs were represented by their molecular structures using Simplified Molecular Input Line Entry System (SMILES) notation. This alphanumeric representation was then converted into a numerical molecular graph representation using RDKit^37^, a cheminformatics package, and Pytorch Geometric^38^. The paper underlines the limitations of molecular fingerprints for representing molecules, emphasizing that structural similarity doesn’t necessarily imply molecular similarity. Drug data in the CTRPv2 dataset was categorized as targeted or untargeted, based on whether they were specifically designed to inhibit genes and gene products involved in cancer-related pathways. Of the 481 drugs in CTRPv2, 31 are classified as targeted, and are FDA-approved.

For drug response data, changes in cellular viability were measured using DNA content, acknowledging that many cell lines respond only partially to the tested drugs. To offer a more intuitive interpretation of the response, this study opted to use the area above the curve (AAC), computed using the PharmacoGx package^39^. Data from the CTRPv2 project was utilized, which ensured consistent cell media and conditions across all tested cell lines to minimize non-drug effects on cell growth. Of the 310,792 samples in CTRPv2, 20,842 were associated with targeted drugs, and 289,950 with untargeted drugs. It was observed that a larger proportion of samples with untargeted drugs had AACs closer to zero, while targeted drugs had a slightly higher median AAC.

### Algorithm

#### Label Distribution Smoothing

Drawing inspiration from classification problems, the Label Distribution Smoothing (LDS) strategy assigns higher weights to data points from less prevalent classes to mitigate dataset imbalance. As DRP falls within the realm of regression tasks, which lack distinct classes, the AAC distribution is divided into bins of width 0.01 within the [0,1] range. Contrary to classification, the proximity of these bins represents their similarity. Using this concept, the LDS technique employs a symmetrical kernel like a Gaussian kernel to smoothen the distribution.

This allows the application of a class prevalence-based weighting scheme to regression-based, continuous data. Subsequently, each bin is assigned a factor reflecting its population size. The inverse of this factor is scaled and multiplied by the loss value during training, compelling the model to emphasize learning from specific data points based on the population of its bin and neighboring bins, addressing dataset imbalance, and improving the model’s generalization ability to data divergent from the average. It should be noted that this method is applied exclusively during training, with sample weights not utilized during inference. See the appendix for details.

#### Autoencoders

To tackle data scarcity and incompleteness issues, we utilized an autoencoder^40^ in our study. This symmetric neural network reduces data dimensionality, minimizing problems associated with high-dimension data and scarcity. Its modularity enables independent training and flexibility in accommodating different data types without requiring extensive re-training. This feature enhances efficiency, reduces interference, and facilitates more straightforward interpretation and biomarker discovery.

#### Graph Neural Networks

In this project, drug molecules are represented as graphs, where atoms are represented as nodes and bonds as edges. Nodes and edges can have properties such as aromaticity, hybridization, bond type, and chirality. GNNs then pass information from each node and its neighbors via its edges iteratively using smaller neural networks to embed either the atoms, the molecule, or both in the latent space. This embedding can then be used in an ANN for learning a specific task. In this project, We use the AttentiveFP GNN model^19^ implementation in the Pytorch Geometric^38^ Python package to extract features from drug molecules for the DRP task. One of the advantages of the AttentiveFP model is its use of the graph attention mechanism^41^, which weighs neighboring nodes based on their relevance to the current node, which has helped it outperform other GNN architectures in most cheminformatic-related tasks. Molecular features used in this project are summarized in **Supplementary Table 5** and **Supplementary Table 6**. See the appendix for more details.

#### Low-rank Multimodal Fusion

Three integration strategies are prevalent in machine learning – early, intermediate, and late integration^42^. This project employs an intermediate integration approach, where an encoder initially embeds each input data type into a latent space. Then these embeddings are combined to create an integrated representation of all inputs. This intermediate method models interactions among the modalities and embedded features, mitigating ambiguities prevalent in unimodal models and enabling better predictions. Multi-linear pooling extends the concept of a two-vector outer product to multiple vectors, resulting in a tensor. The tensor fusion method further models the combinations of each modality^43^. A low-rank approximation of this approach mitigates computational burden while allowing for context-specific weights for each modality, addressing issues inherent in concatenation and weighted sum methods^20^. See the appendix for more details.

### Hyper-parameter optimization

In this work, We use the HyperOpt^44^ algorithm from the Ray^45^ package’s Tune^46^ library, which can run trials in parallel. While searching for optimal hyperparameters is a computationally intensive process, it ensures the most effective use of available data. In this project, both training and architectural hyper-parameters were adjusted. To use computational resources effectively, we used the same activation functions, batch normalization, and dropout parameters across the model instead of tailoring them to individual layers.

Autoencoder modules underwent hyperparameter optimization first, given their preliminary training on their respective data sets. After identifying optimal hyperparameters for each module, the encoder subnetworks are integrated with the drug encoder to the DRP module. A subsequent hyperparameter search is conducted only for the DRP module and the tensor fusion mechanism. The pre-optimized encoders undergo additional training in the DRP context (**Figure 2**).

Hyperparameter optimization trials used grouped and stratified cross-validation for the complete model, leading to average validation losses across the K folds serving as a final score for each trial. This approach may lead to minor information leakage between cross-validation and hyperparameter optimization processes. A nested cross-validation scheme can mitigate this issue, but due to computational demands and time constraints, this method was not implemented; instead, trial numbers were capped at 40 per model type. See the appendix for details.

### Stratified Group K-fold Cross-validation

The predictive performance of the model was evaluated using k-fold cross-validation^47^. However, this non-exhaustive method might lead to unequal distribution of different cancer types in training and validation sets, potentially reducing the evaluation’s representativeness of real-world performance. To mitigate this, stratification and strict subsetting were employed to maintain uniform distribution of data points and assess model generalization. Stratification ensures equal proportions of cell lines based on their lineage in each bin or fold, avoiding bias towards specific lineages or cell line sets. Strict subsetting, contrasted with lenient subsetting, ensures the model only encounters previously unseen cell lines, drugs, or both in the validation set, thereby assessing the model’s response to novel scenarios.

Stratified Group K-fold from the scikit-learn python package^48^ was used for this purpose. The combination of stratification and grouping enabled a more reliable evaluation of the model’s ability to generalize to new tasks (**Supplementary** Figure 9). See the appendix for more details.

### Interpretation using Integrated Gradients

There are different classes of ML interpretation techniques based on various criteria. Some models, such as shallow decision trees and sparse linear models, are intrinsically interpretable by analyzing internal model parameters. Other models require post-hoc interpretation, which is done by analyzing the model after training^49^. Furthermore, interpretation can be model-specific or model-agnostic, and some methods also seek to explain a complex model through a simpler, more interpretable conjugate model. A few well-known interpretation methods are provided in the Captum package for Pytorch^50^, which we have used. Specifically, For this task, we used the Integrated Gradients technique^51^, which is more suitable for neural networks. We then applied this interpretation method to identify important biological features and drug characteristics.

