## Supplementary Tables and Figures for "Drug Response Prediction and Biomarker Discovery Using Multi-Modal Deep Learning"

**Supplementary Table 1. Targeted drugs in CTRPv2 by cancer type.**

| Cancer Type | Targeted Drugs |
|---|---|
| **Leukemia** | Idelalisib, Venetoclax, Tretinoin, Dasatinib, Ibrutinib, Nilotinib, Bosutinib |
| **Breast Cancer** | Olaparib, Tamoxifen, Lapatinib, Fulvestrant, Alpelisib |
| **Lung Cancer** | Crizotinib, Erlotinib, Gefitinib, Dabrafenib |
| **Colorectal Cancer** | Regorafenib |
| **Lymphoma** | Bortezomib, Bexarotene, Belinostat, Vorinostat |
| **Thyroid Cancer** | Cabozantinib, Vandetanib |
| **Skin Cancer** | Sonidegib, Trametinib, Vemurafenib |
| **Kidney Cancer** | Axitinib, Temsirolimus, Sorafenib |
| **Soft Tissue Sarcoma** | Pazopanib |
| **Pancreatic Cancer** | Sunitinib |

**Supplementary Table 2. Databases relevant to the drug response prediction task used in this project.**

| Resource | Data | Description |
|---|---|---|
| DepMap (21Q2) | **Mutation:** 18,802 genes, 1,734 cell lines (WGS)<br>**Copy Number:** 27,561 gene overlaps, 1,742 cell lines (Mostly WGS) | 1,814 Cell Lines Overall (Incomplete Data) |

| | | |
|---|---|---|
| | **Gene Expression:** 19,176 genes, 1,378 cell lines (RNA-Seq)<br>**Protein Quantification: (**up to) 12,399 proteins, 5,153 shared across all cell lines, 378 cell lines<br>**Histone Profiling:** 42 histone modification sites, 896 cell lines<br>**miRNA Expression:** 734 miRNAs, 953 cell lines<br>**Metabolomics:** 225 metabolites, 927 cell lines<br>**RPPA:** 214 proteins, 898 cell lines | |
| CTRPv2 | **Compound**: Structure, Synonyms, Target, Concentrations<br>**Dose-Response:** 310,792 Dose-Response Values | 860 cell lines (rows) by 481 drugs (columns) (incomplete) |
| ChEMBL[58,59] (version 27) | **Compound**: SMILES representation of drug-like molecules | 1,941,411 Compounds |

**Supplementary Table 3. Best bimodal models and their RMSE losses by omic type, splitting method, and drug type in the AAC ≥ 0.7 range.**

| Omic Type | Split By Both Cell Line & Drug Scaffold | | Split By Cell Line | | Split By Drug Scaffold | |
|---|---|---|---|---|---|---|
| | Targeted Drug | Untargeted Drug | Targeted Drug | Untargeted Drug | Targeted Drug | Untargeted Drug |
| MUT | LDS+LMF<br>0.599 | LMF<br>0.28 | LDS+LMF<br>0.566 | LMF<br>0.334 | LDS+LMF+GNN<br>0.547 | LDS+LMF+GNN<br>0.221 |

| Omic Type | Split By Both Cell Line & Drug Scaffold | | Split By Cell Line | | Split By Drug Scaffold | |
|---|---|---|---|---|---|---|
| | Targeted Drug | Untargeted Drug | Targeted Drug | Untargeted Drug | Targeted Drug | Untargeted Drug |
| CNV | LMF+GNN<br>0.448 | LMF<br>0.209 | LMF<br>0.431 | LDS+LMF+GNN<br>0.21 | LDS+LMF+GNN<br>0.435 | LDS+LMF+GNN<br>0.195 |
| EXP | LMF<br>0.456 | LMF<br>0.19 | LMF<br>0.451 | LMF<br>0.185 | LDS+LMF+GNN<br>0.403 | LDS+LMF+GNN<br>0.166 |
| PROT | LMF+GNN<br>0.48 | LDS+LMF+GNN<br>0.211 | LMF+GNN<br>0.454 | GNN<br>0.208 | LDS+LMF+GNN<br>0.246 | LDS+LMF+GNN<br>0.13 |
| MIRNA | LMF+GNN<br>0.542 | LMF<br>0.228 | LMF<br>0.435 | LMF<br>0.213 | LMF<br>0.49 | LDS+LMF+GNN<br>0.21 |
| METAB | LMF+GNN<br>0.503 | LDS+LMF<br>0.219 | LDS+LMF+GNN<br>0.576 | LMF<br>0.232 | LDS+LMF+GNN<br>0.542 | LDS+LMF+GNN<br>0.209 |
| HIST | LDS+LMF+GNN<br>0.579 | LDS+LMF+GNN<br>0.245 | GNN<br>0.572 | GNN<br>0.259 | Sum<br>0.569 | LDS+LMF+GNN<br>0.235 |
| RPPA | GNN<br>0.524 | LMF<br>0.204 | Sum<br>0.484 | LMF<br>0.209 | LDS+LMF+GNN<br>0.489 | LDS+LMF+GNN<br>0.214 |

**Supplementary Table 4. Summary of top repurposable drug candidates (Empirical AAC ≥ Highest Prescribed AAC + 0.2) paired with the model that can predict the empirical AAC most accurately (MAE ≤ 0.2). Note that the empirical cell line is novel to the model.**

| Empirical | | | | Prescribed | | | Top Model: Data Type(s) & Prediction |
|---|---|---|---|---|---|---|---|
| Primary Disease | Lineage Subtype | Cell Line | AAC | Highest AAC | Cancer | Drug | |
| Brain Cancer | Glioma | AM38 | 0.684 | 0.367 | Breast Cancer | Alpelisib | HIST + RPPA<br>0.73 |

| Empirical | | | | Prescribed | | | Top Model: Data Type(s) & Prediction |
|---|---|---|---|---|---|---|---|
| **Primary Disease** | **Lineage Subtype** | **Cell Line** | **AAC** | **Highest AAC** | **Cancer** | **Drug** | |
| Breast Cancer | Breast Carcinoma | DU4475 | 0.953 | 0.293 | Lung Cancer | Dabrafenib | MUT + EXP 0.777 |
| | | HCC1569 | 0.567 | 0.299 | Kidney Cancer | Temsirolimus | METAB + RPPA 0.598 |
| | Breast Ductal Carcinoma | HCC1954 | 0.631 | | | | MIRNA + HIST 0.612 |
| Colon/Colorectal Cancer | Colorectal Adenocarcinoma | NCIH716 | 0.481 | 0.253 | Sarcoma | Pazopanib | MUT + PROT 0.38 |
| Endometrial/Uterine Cancer | Endometrial Adenocarcinoma | JHUEM2 | 0.564 | 0.299 | Kidney Cancer | Temsirolimus | MIRNA + HIST 0.559 |
| Head and Neck Cancer | Upper Aerodigestive Squamous | YD10B | 0.508 | 0.304 | Thyroid Cancer | Vandetanib | MUT + EXP 0.506 |
| Leukemia | ALL | KE37 | 0.618 | 0.299 | Kidney Cancer | Temsirolimus | MUT + HIST 0.623 |
| | | MOLT16 | 0.603 | | | | MUT + EXP 0.605 |

| Empirical | | | | Prescribed | | | Top Model: Data Type(s) & Prediction |
|---|---|---|---|---|---|---|---|
| Primary Disease | Lineage Subtype | Cell Line | AAC | Highest AAC | Cancer | Drug | |
| | | SEM | 0.657 | 0.282 | Pancreatic Cancer | Sunitinib | PROT + MIRNA 0.661 |
| | AML | EOL1 | 0.992 | 0.379 | Colon/Colorectal Cancer | Regorafenib | MUT + PROT 0.991 |
| | | | 0.991 | 0.495 | Thyroid Cancer | Cabozantinib | PROT + MIRNA 0.928 |
| | | | 0.988 | 0.385 | Kidney Cancer | Sorafenib | PROT + MIRNA 0.99 |
| | | | 0.981 | 0.554 | | Axitinib | PROT + RPPA 0.913 |
| | | | 0.967 | 0.253 | Sarcoma | Pazopanib | MUT + PROT 0.816 |
| | | | 0.952 | 0.282 | Pancreatic Cancer | Sunitinib | MUT + PROT 0.941 |
| | | | 0.583 | 0.304 | Thyroid Cancer | Vandetanib | MUT + PROT 0.569 |

| Empirical | | | | Prescribed | | | Top Model: Data Type(s) & Prediction |
|---|---|---|---|---|---|---|---|
| **Primary Disease** | **Lineage Subtype** | **Cell Line** | **AAC** | **Highest AAC** | **Cancer** | **Drug** | |
| | | KASUMI1 | 0.659 | 0.282 | Pancreatic Cancer | Sunitinib | CNV + EXP 0.695 |
| | | | 0.647 | 0.385 | Kidney Cancer | Sorafenib | PROT + RPPA 0.674 |
| | | | 0.576 | 0.253 | Sarcoma | Pazopanib | PROT + HIST 0.547 |
| | | MOLM13 | 0.855 | 0.495 | Thyroid Cancer | Cabozantinib | MUT + PROT 0.843 |
| | | | 0.839 | 0.282 | Pancreatic Cancer | Sunitinib | MUT + RPPA 0.841 |
| | | | 0.793 | 0.385 | Kidney Cancer | Sorafenib | MUT + RPPA 0.802 |
| | | | 0.504 | 0.299 | | Temsirolimus | PROT + MIRNA 0.505 |
| | | MOLM16 | 0.704 | 0.495 | Thyroid Cancer | Cabozantinib | CNV + MIRNA 0.62 |

| Empirical | | | | Prescribed | | | Top Model: Data Type(s) & Prediction |
|---|---|---|---|---|---|---|---|
| **Primary Disease** | **Lineage Subtype** | **Cell Line** | **AAC** | **Highest AAC** | **Cancer** | **Drug** | |
| | | MONOMAC1 | 0.541 | 0.299 | Kidney Cancer | Temsirolimus | EXP + MIRNA 0.542 |
| | | MV411 | 0.886 | 0.385 | Kidney Cancer | Sorafenib | METAB + RPPA 0.838 |
| | | MV411 | 0.829 | 0.282 | Pancreatic Cancer | Sunitinib | CNV + EXP 0.842 |
| | | MV411 | 0.655 | 0.299 | Kidney Cancer | Temsirolimus | METAB 0.644 |
| | | THP1 | 0.515 | 0.299 | Kidney Cancer | Temsirolimus | MUT + MIRNA 0.515 |
| | CLL | JVM3 | 0.745 | 0.299 | Kidney Cancer | Temsirolimus | CNV + MIRNA 0.732 |
| Lung Cancer | NSCLC | CALU3 | 0.512 | 0.299 | Kidney Cancer | Temsirolimus | MIRNA + HIST 0.51 |
| Lung Cancer | NSCLC | EBC1 | 0.543 | 0.299 | Kidney Cancer | Temsirolimus | MUT + CNV 0.55 |

| Empirical | | | | Prescribed | | | Top Model: Data Type(s) & Prediction |
|---|---|---|---|---|---|---|---|
| **Primary Disease** | **Lineage Subtype** | **Cell Line** | **AAC** | **Highest AAC** | **Cancer** | **Drug** | |
| | | HCC827 | 0.520 | 0.304 | Thyroid Cancer | Vandetanib | EXP + RPPA + HIST + PROT 0.52 |
| | | NCIH1703 | 0.596 | 0.282 | Pancreatic Cancer | Sunitinib | CNV + PROT 0.644 |
| | | | 0.476 | 0.253 | Sarcoma | Pazopanib | MUT + PROT 0.47 |
| | | PC14 | 0.597 | 0.304 | Thyroid Cancer | Vandetanib | MUT + EXP 0.602 |
| | SCLC | NCIH211 | 0.550 | 0.299 | Kidney Cancer | Temsirolimus | MUT + METAB 0.544 |
| | Mesothelioma | MSTO211H | 0.621 | | | | CNV + MIRNA 0.614 |
| | | NCIH2452 | 0.580 | | | | MIRNA + METAB 0.553 |

| Empirical | | | | Prescribed | | | Top Model: Data Type(s) & Prediction |
|---|---|---|---|---|---|---|---|
| Primary Disease | Lineage Subtype | Cell Line | AAC | Highest AAC | Cancer | Drug | |
| Lymphoma | Hodgkin Lymphoma | L540 | 0.545 | | | | MUT + MIRNA 0.546 |
| | Non-Hodgkin Lymphoma | HT | 0.616 | | | | CNV + MIRNA 0.621 |
| | | KARPAS299 | 0.596 | | | | MIRNA + METAB 0.594 |
| | | KIJK | 0.659 | | | | MUT + EXP 0.658 |
| | | NUDUL1 | 0.629 | | | | MUT + EXP 0.63 |
| | | RI1 | 0.704 | | | | CNV + EXP 0.706 |
| | | | 0.699 | 0.496 | Leukemia | Ibrutinib | CNV + EXP 0.595 |
| | | SR786 | 0.612 | 0.299 | Kidney Cancer | Temsirolimus | MUT + CNV 0.614 |

| Empirical | | | | Prescribed | | | Top Model: Data Type(s) & Prediction |
|---|---|---|---|---|---|---|---|
| **Primary Disease** | **Lineage Subtype** | **Cell Line** | **AAC** | **Highest AAC** | **Cancer** | **Drug** | |
| | | SUDHL4 | 0.540 | | | | CNV + PROT 0.544 |
| | | SUDHL6 | 0.746 | | | | MIRNA + RPPA 0.746 |
| | | | 0.725 | 0.477 | Leukemia | Idelalisib | CNV + EXP 0.538 |
| | | SUDHL8 | 0.617 | 0.299 | Kidney Cancer | Temsirolimus | EXP + METAB 0.611 |
| | | WSUDLCL2 | 0.676 | | | | EXP + MIRNA 0.669 |
| Myeloma | Multiple Myeloma | KE97 | 0.589 | | | | MIRNA + RPPA 0.593 |
| | | KMS11 | 0.515 | | | | EXP + RPPA + HIST + PROT 0.518 |

| Empirical | | | | Prescribed | | | Top Model: Data Type(s) & Prediction |
|---|---|---|---|---|---|---|---|
| Primary Disease | Lineage Subtype | Cell Line | AAC | Highest AAC | Cancer | Drug | |
| | | KMS12BM | 0.539 | | | | EXP + PROT 0.537 |
| | | KMS21BM | 0.669 | | | | CNV 0.686 |
| | | KMS26 | 0.837 | | | | MIRNA + METAB 0.778 |
| | | KMS34 | 0.824 | | | | MUT + MIRNA 0.818 |
| | | L363 | 0.650 | | | | MIRNA + HIST 0.645 |
| | | MM1S | 0.523 | | | | MUT + EXP 0.522 |
| Ovarian Cancer | Ovary Adenocarcinoma | A2780 | 0.622 | | | | CNV + EXP 0.621 |
| | | OVK18 | 0.537 | | | | CNV + RPPA 0.537 |

<table>
<tr><th colspan="4">Empirical</th><th colspan="3">Prescribed</th><th rowspan="2">Top Model: Data Type(s) & Prediction</th></tr>
<tr><th>Primary Disease</th><th>Lineage Subtype</th><th>Cell Line</th><th>AAC</th><th>Highest AAC</th><th>Cancer</th><th>Drug</th></tr>
<tr><td>Pancreatic Cancer</td><td>Exocrine</td><td>PATU8988S</td><td>0.510</td><td>0.304</td><td>Thyroid Cancer</td><td>Vandetanib</td><td>MIRNA + METAB 0.313</td></tr>
<tr><td rowspan="2">Rhabdoid</td><td rowspan="2">Malignant Rhabdoid Tumor</td><td rowspan="2">A204</td><td>0.597</td><td>0.282</td><td>Pancreatic Cancer</td><td>Sunitinib</td><td>CNV + PROT 0.609</td></tr>
<tr><td>0.473</td><td>0.253</td><td>Sarcoma</td><td>Pazopanib</td><td>CNV + EXP 0.424</td></tr>
<tr><td rowspan="5">Skin Cancer</td><td rowspan="5">Melanoma</td><td>COLO829</td><td>0.520</td><td rowspan="5">0.293</td><td rowspan="5">Lung Cancer</td><td rowspan="5">Dabrafenib</td><td>PROT + RPPA 0.533</td></tr>
<tr><td>HT144</td><td>0.521</td><td>CNV + EXP 0.493</td></tr>
<tr><td>SH4</td><td>0.674</td><td>MUT + EXP 0.604</td></tr>
<tr><td>SKMEL24</td><td>0.622</td><td>CNV + EXP 0.565</td></tr>
<tr><td>UACC257</td><td>0.613</td><td>CNV + EXP 0.624</td></tr>
</table>

| Empirical | | | | Prescribed | | | Top Model: Data Type(s) & Prediction |
|---|---|---|---|---|---|---|---|
| Primary Disease | Lineage Subtype | Cell Line | AAC | Highest AAC | Cancer | Drug | |
| | | WM115 | 0.755 | | | | EXP + PROT 0.793 |
| | | WM2664 | 0.575 | | | | MUT + MIRNA 0.572 |
| | | WM983B | 0.578 | | | | CNV + EXP 0.572 |
| Thyroid Cancer | Thyroid Squamous | SW579 | 0.531 | 0.282 | Pancreatic Cancer | Sunitinib | CNV + EXP 0.54 |
| | | | 0.469 | 0.253 | Sarcoma | Pazopanib | MUT + EXP 0.331 |

**Supplementary Table 5. Atomic (node) features for the AttentiveFP model.**

| Atom feature | Size | Description |
|---|---|---|
| Atom symbol | 13 | ['As', 'B', 'Br', 'C', 'Cl', 'F', 'We', 'N', 'O', 'P', 'Pt', 'S', 'other'] (one-hot) |
| Degree | 6 | Number of covalent bonds [0, 1, 2, 3, 4, 5] (one-hot) |
| Formal charge | 1 | Electrical charge (integer) |
| Radical electrons | 1 | Number of radical electrons (integer) |

| | | |
|---|---|---|
| Hybridization | 7 | [sp, sp2, sp3, sp3d, sp3d2, unspecified, other] (one-hot) |
| Aromaticity | 1 | whether the atom is part of an aromatic system [0/1] (one-hot) |
| Hydrogens | 5 | number of connected hydrogens [0,1,2,3,4] (one-hot) |
| Chirality | 1 | whether the atom is chiral center [0/1] (one-hot) |
| Chirality type | 2 | [R, S] (one-hot) |

**Supplementary Table 6. Bond features for the AttentiveFP model.**

| Bond feature | Size | Description |
|---|---|---|
| Bond type | 4 | [single, double, triple, aromatic] (one-hot) |
| Conjugation | 1 | whether the bond is conjugated [0/1] (one-hot) |
| Ring | 1 | whether the bond is in a ring [0/1] (one-hot) |
| Stereo | 4 | [StereoNone, StereoAny, StereoZ, StereoE] (one-hot) |

**Supplementary Table 7. Proportion of bimodal validation samples with predictions within 0.2 of the ground truth AAC target.**

| Data Type(s) | Configuration | Target Below 0.7 | | Target Above 0.7 | |
|---|---|---|---|---|---|
| | | Untargeted Drug | Targeted Drug | Untargeted Drug | Targeted Drug |
| MUT | Baseline | 0.9296 | 0.9156 | 0.5691 | 0.0000 |
| | Trifecta | 0.9270 | 0.9158 | 0.6169 | 0.0222 |
| CNV | Baseline | 0.9692 | 0.9657 | 0.6912 | 0.0000 |

| Data Type(s) | Configuration | Target Below 0.7 | | Target Above 0.7 | |
|---|---|---|---|---|---|
| | | Untargeted Drug | Targeted Drug | Untargeted Drug | Targeted Drug |
| | Trifecta | 0.9695 | 0.9639 | 0.7625 | 0.1111 |
| EXP | Baseline | 0.9723 | 0.9678 | 0.7566 | 0.0444 |
| | Trifecta | 0.9766 | 0.9731 | 0.7816 | 0.1556 |
| PROT | Baseline | 0.9688 | 0.9612 | 0.7287 | 0.0000 |
| | Trifecta | 0.9711 | 0.9672 | 0.7353 | 0.0889 |
| MIRNA | Baseline | 0.9684 | 0.9602 | 0.6949 | 0.0000 |
| | Trifecta | 0.9661 | 0.9581 | 0.7228 | 0.0222 |
| METAB | Baseline | 0.9667 | 0.9611 | 0.7015 | 0.0000 |
| | Trifecta | 0.9625 | 0.9525 | 0.7000 | 0.0222 |
| HIST | Baseline | 0.9638 | 0.9541 | 0.6125 | 0.0000 |
| | Trifecta | 0.9628 | 0.9547 | 0.6478 | 0.0000 |
| RPPA | Baseline | 0.9718 | 0.9634 | 0.7265 | 0.0000 |
| | Trifecta | 0.9640 | 0.9582 | 0.7169 | 0.0000 |

**Supplementary Table 8. Proportion of trimodal validation samples with predictions within 0.2 of the ground truth AAC target.**

| Data Type(s) | Configuration | Target Below 0.7 | | Target Above 0.7 | |
|---|---|---|---|---|---|
| | | Untargeted Drug | Targeted Drug | Untargeted Drug | Targeted Drug |
| CNV + EXP | Baseline | 0.9703 | 0.9658 | 0.7434 | 0.0222 |
| | Trifecta | 0.9930 | 0.9924 | 0.9375 | 0.7778 |
| CNV + HIST | Baseline | 0.9704 | 0.9669 | 0.7316 | 0.0222 |
| | Trifecta | 0.9794 | 0.9734 | 0.8346 | 0.2444 |
| CNV + METAB | Baseline | 0.9698 | 0.9622 | 0.7419 | 0.0889 |
| | Trifecta | 0.9825 | 0.9797 | 0.8625 | 0.3333 |
| CNV + MIRNA | Baseline | 0.9712 | 0.9656 | 0.7176 | 0.0667 |
| | Trifecta | 0.9852 | 0.9833 | 0.8669 | 0.4667 |
| CNV + PROT | Baseline | 0.9675 | 0.9601 | 0.6515 | 0.0000 |
| | Trifecta | 0.9915 | 0.9909 | 0.9331 | 0.6222 |
| CNV + RPPA | Baseline | 0.9698 | 0.9657 | 0.6978 | 0.0000 |
| | Trifecta | 0.9834 | 0.9818 | 0.8779 | 0.5333 |
| EXP + HIST | Baseline | 0.9729 | 0.9697 | 0.7787 | 0.0889 |
| | Trifecta | 0.9807 | 0.9794 | 0.8331 | 0.2000 |

| Data Type(s) | Configuration | Target Below 0.7 | | Target Above 0.7 | |
|---|---|---|---|---|---|
| | | Untargeted Drug | Targeted Drug | Untargeted Drug | Targeted Drug |
| EXP + METAB | Baseline | 0.9722 | 0.9672 | 0.7647 | 0.0889 |
| | Trifecta | 0.9851 | 0.9837 | 0.8824 | 0.4222 |
| EXP + MIRNA | Baseline | 0.9716 | 0.9668 | 0.7684 | 0.0889 |
| | Trifecta | 0.9813 | 0.9776 | 0.8228 | 0.1556 |
| EXP + PROT | Baseline | 0.9644 | 0.9513 | 0.6912 | 0.0222 |
| | Trifecta | 0.9895 | 0.9873 | 0.9051 | 0.5778 |
| EXP + RPPA | Baseline | 0.9713 | 0.9662 | 0.7684 | 0.1556 |
| | Trifecta | 0.9870 | 0.9883 | 0.8831 | 0.4444 |
| HIST + RPPA | Baseline | 0.9676 | 0.9657 | 0.7074 | 0.0444 |
| | Trifecta | 0.9829 | 0.9766 | 0.8663 | 0.3095 |
| METAB + HIST | Baseline | 0.9691 | 0.9610 | 0.6949 | 0.0000 |
| | Trifecta | 0.9698 | 0.9618 | 0.7500 | 0.0444 |
| METAB + RPPA | Baseline | 0.9713 | 0.9661 | 0.7368 | 0.0222 |
| | Trifecta | 0.9721 | 0.9705 | 0.7963 | 0.3111 |
| MIRNA + HIST | Baseline | 0.9685 | 0.9642 | 0.6831 | 0.0000 |
| | Trifecta | 0.9826 | 0.9756 | 0.8667 | 0.2500 |

| Data Type(s) | Configuration | Target Below 0.7 | | Target Above 0.7 | |
|---|---|---|---|---|---|
| | | Untargeted Drug | Targeted Drug | Untargeted Drug | Targeted Drug |
| MIRNA + METAB | Baseline | 0.9699 | 0.9642 | 0.7037 | 0.0000 |
| | Trifecta | 0.9889 | 0.9855 | 0.9272 | 0.6000 |
| MIRNA + RPPA | Baseline | 0.9696 | 0.9642 | 0.7140 | 0.0222 |
| | Trifecta | 0.9868 | 0.9843 | 0.8918 | 0.5946 |
| MUT + CNV | Baseline | 0.8891 | 0.8778 | 0.6441 | 0.0222 |
| | Trifecta | 0.9878 | 0.9847 | 0.8934 | 0.5556 |
| MUT + EXP | Baseline | 0.9233 | 0.9163 | 0.6463 | 0.0222 |
| | Trifecta | 0.9923 | 0.9919 | 0.9257 | 0.6889 |
| MUT + HIST | Baseline | 0.9417 | 0.9263 | 0.6118 | 0.0000 |
| | Trifecta | 0.9808 | 0.9777 | 0.8250 | 0.2667 |
| MUT + METAB | Baseline | 0.9403 | 0.9274 | 0.6603 | 0.0000 |
| | Trifecta | 0.9852 | 0.9790 | 0.8581 | 0.3778 |
| MUT + MIRNA | Baseline | 0.9272 | 0.9135 | 0.6110 | 0.0000 |
| | Trifecta | 0.9861 | 0.9826 | 0.8882 | 0.4889 |
| MUT + PROT | Baseline | 0.9239 | 0.9066 | 0.5809 | 0.0000 |
| | Trifecta | 0.9908 | 0.9894 | 0.9007 | 0.6444 |

| Data Type(s) | Configuration | Target Below 0.7 | | Target Above 0.7 | |
|---|---|---|---|---|---|
| | | Untargeted Drug | Targeted Drug | Untargeted Drug | Targeted Drug |
| MUT + RPPA | Baseline | 0.9269 | 0.9126 | 0.5941 | 0.0000 |
| | Trifecta | 0.9849 | 0.9846 | 0.8691 | 0.5778 |
| PROT + HIST | Baseline | 0.9687 | 0.9580 | 0.7081 | 0.0000 |
| | Trifecta | 0.9869 | 0.9836 | 0.8963 | 0.4000 |
| PROT + METAB | Baseline | 0.9671 | 0.9575 | 0.6721 | 0.0000 |
| | Trifecta | 0.9901 | 0.9877 | 0.9235 | 0.5556 |
| PROT + MIRNA | Baseline | 0.9655 | 0.9534 | 0.6551 | 0.0222 |
| | Trifecta | 0.9906 | 0.9899 | 0.9353 | 0.7111 |
| PROT + RPPA | Baseline | 0.9683 | 0.9603 | 0.7221 | 0.0000 |
| | Trifecta | 0.9900 | 0.9881 | 0.9324 | 0.7778 |

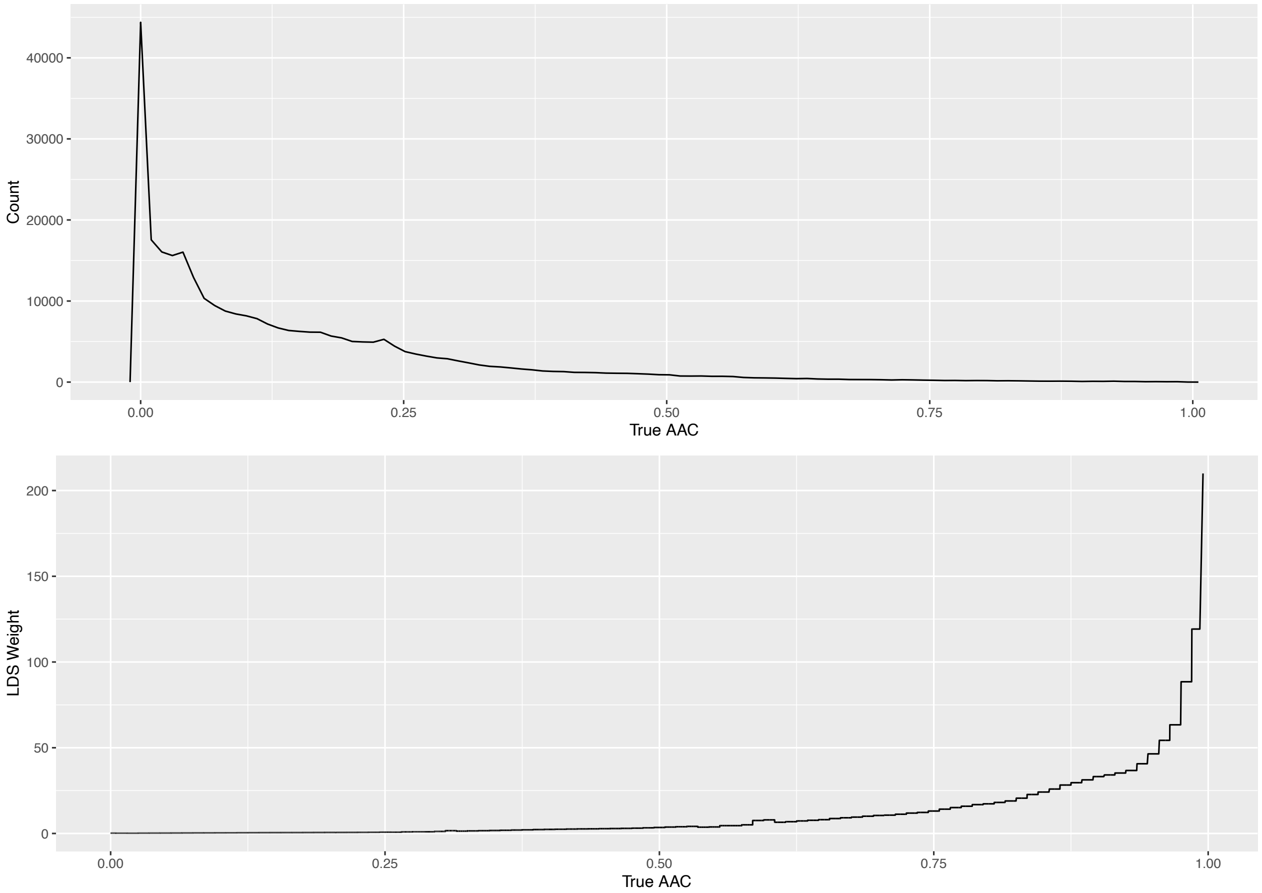


**Supplementary Figure 1. Distribution of AAC (top) vs LDS-calculated weights (bottom) in the CTRPv2 dataset.** LDS assigns higher weights to less frequently observed samples, which in the case of CTRPv2 also have higher AACs. The LDS weight distribution is roughly the inverse of the AAC distribution.

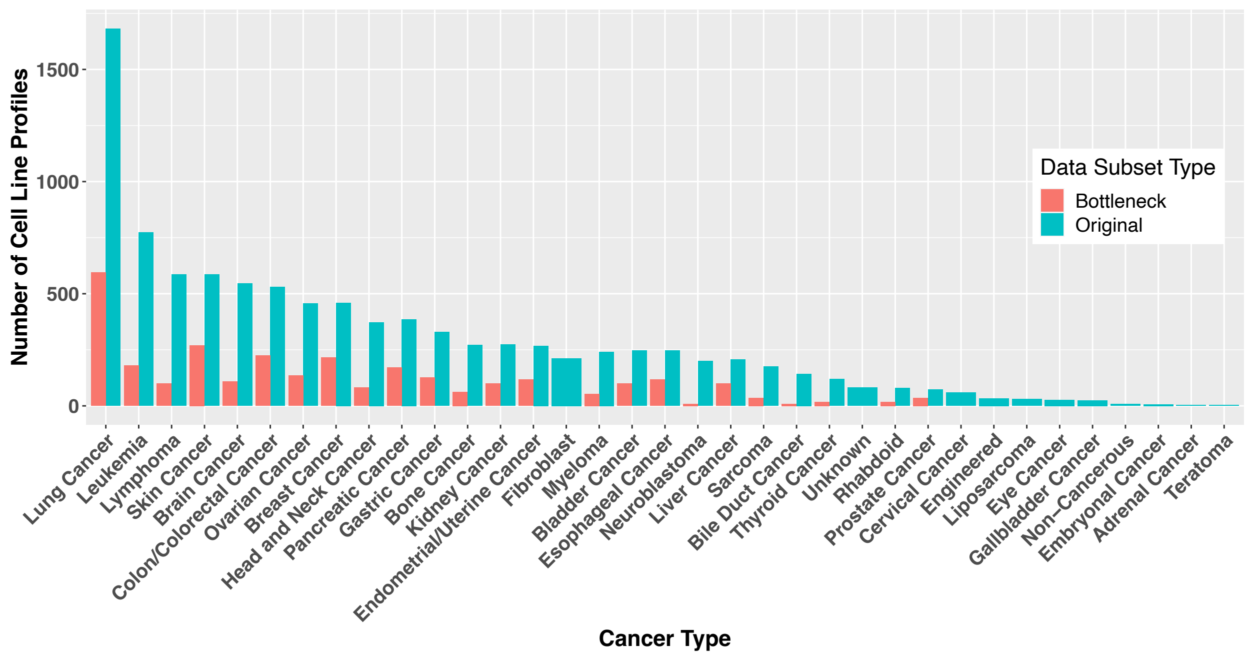


**Supplementary Figure 2. The subset of complete samples by cancer types in DepMap.** Some cancer types do not have cell lines that are characterized with all eight omic profiling technologies. Because of this, they are not represented in the bottlenecked dataset.

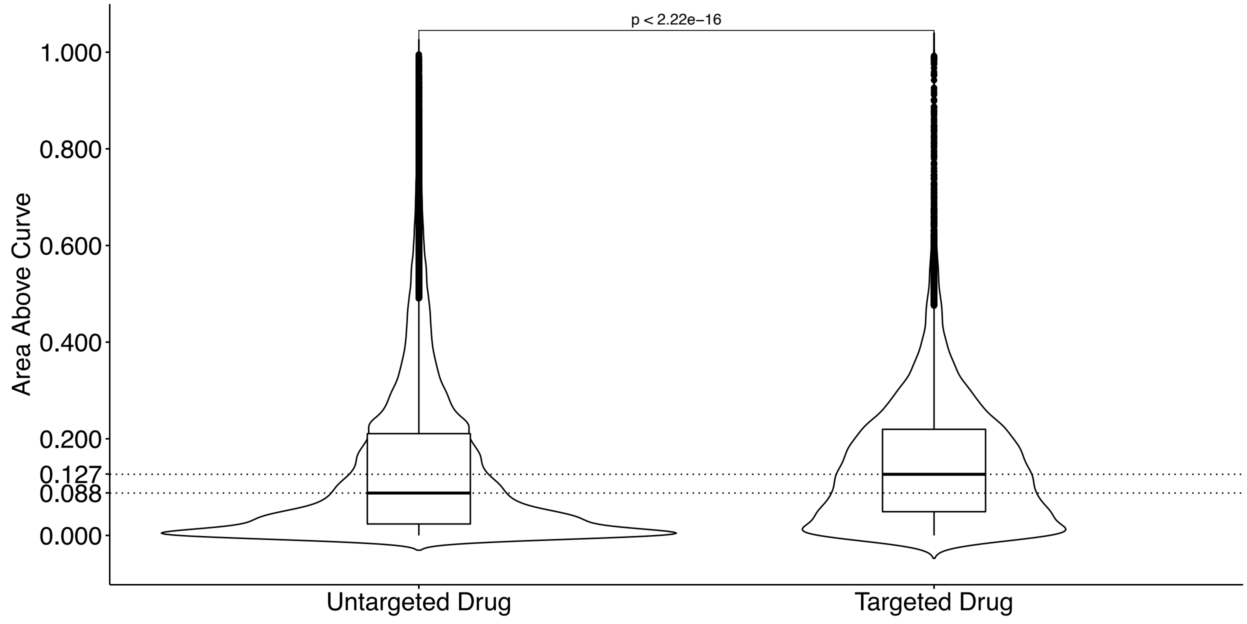


**Supplementary Figure 3. Violin plot and boxplot comparison of AAC distributions of samples with targeted (n=20,842) and untargeted (n=289,950) drugs.** The violin plot depicts the density at each AAC, whereas the boxplot indicates the lower quartile, median and upper quartile of the AAC distributions for targeted and untargeted drugs. Dotted lines indicate the medians of each distribution. The median AAC for targeted drugs is 0.127, which is higher than the median AAC for untargeted drugs at 0.088 (p-value < $2.22 \times 10^{-16}$, Mann-Whitney U Test).

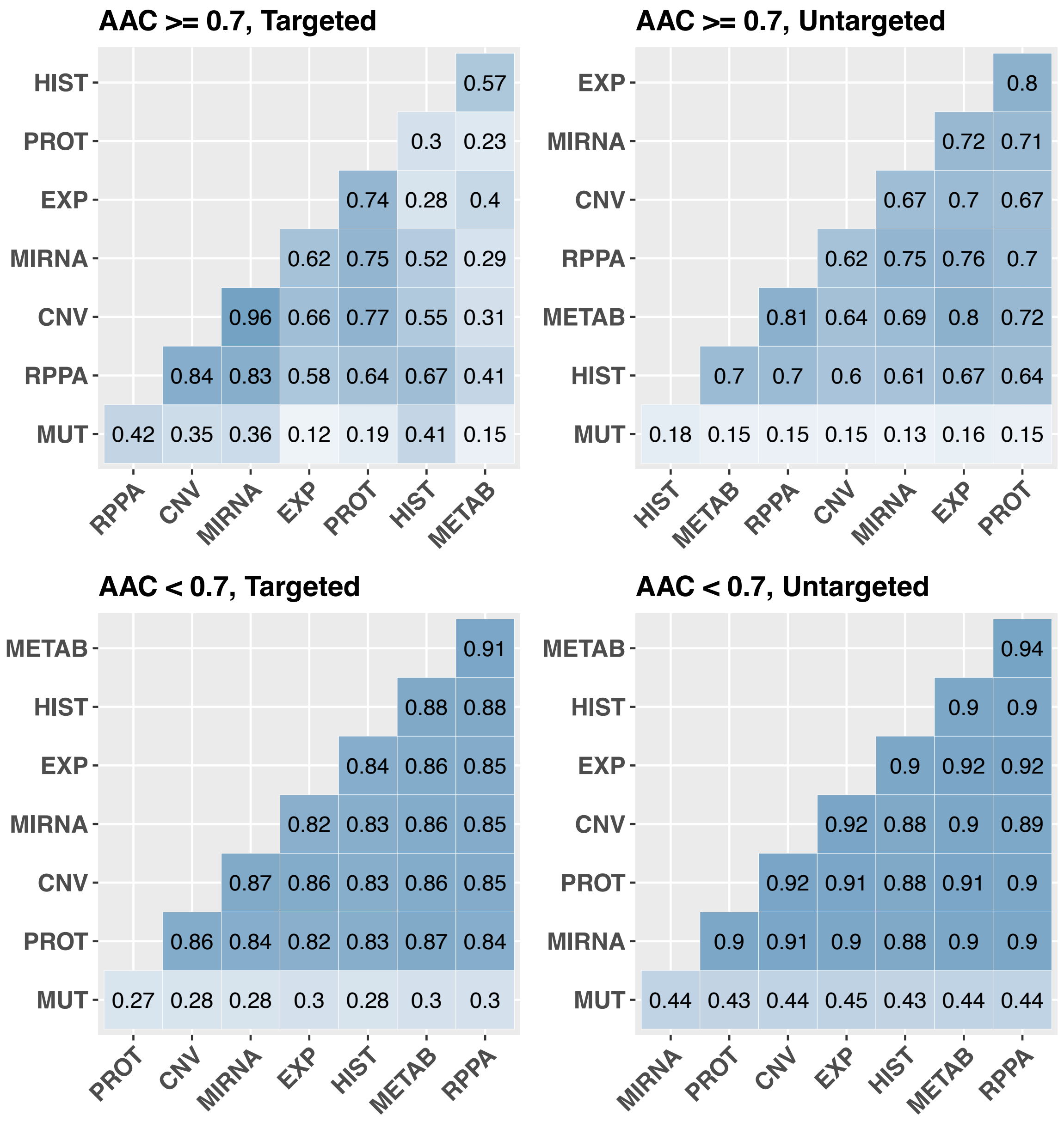


**Supplementary Figure 4. R2 measure of agreement between predictions of bimodal baseline models in distinct AAC range and drug type subsets.** The agreement between different models in the higher AAC ranges, especially in samples with targeted drugs, is much less than in other range and drug type. This indicates the certain omic profiling data may be more useful for specific cell line and drug combinations, and for the mechanisms of action of targeted drugs.

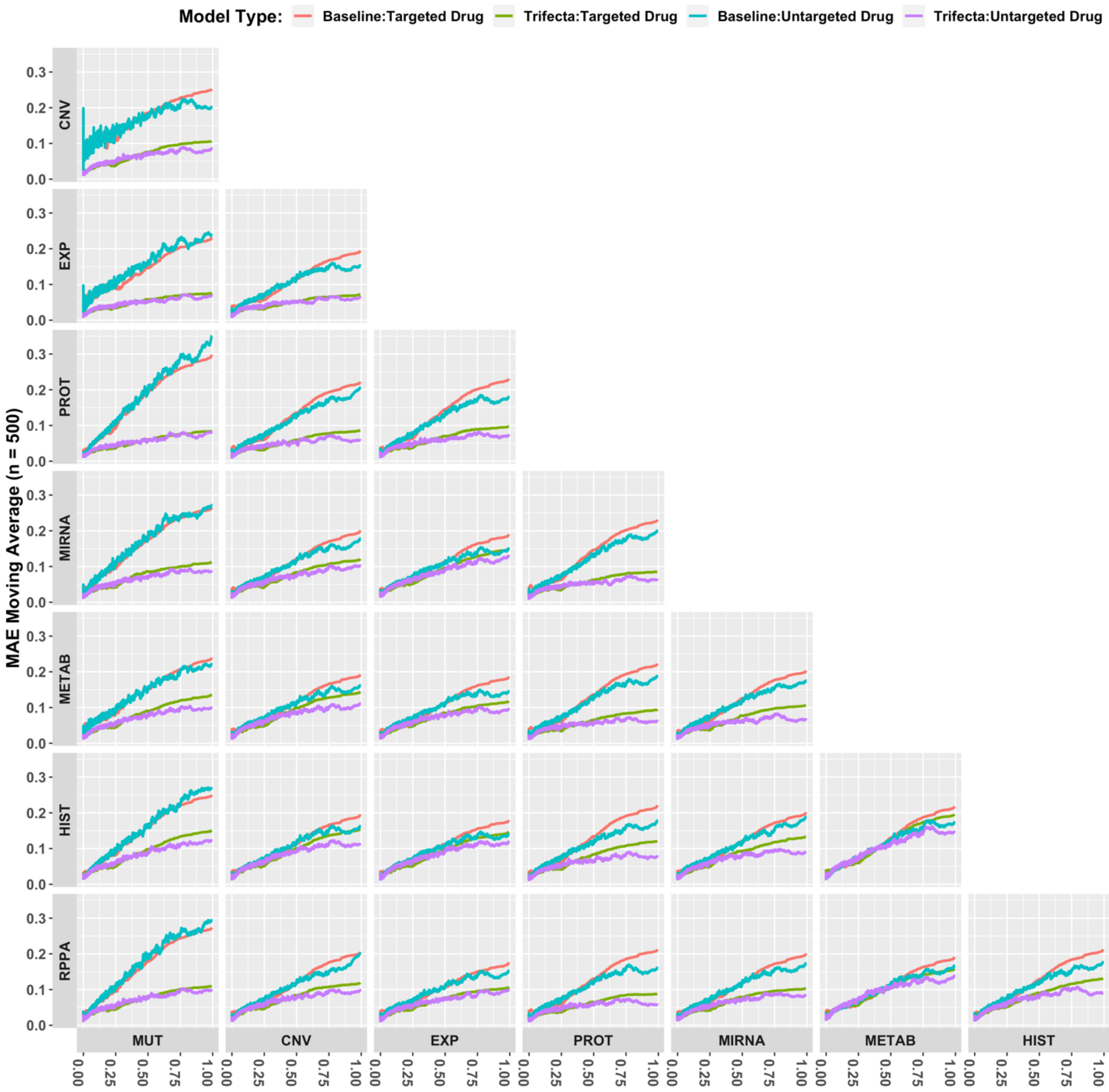


**Supplementary Figure 5. Comparison of trimodal baseline vs. trifecta models.** Trailing moving average of MAE losses in baseline and trifecta models by drug type in each sub-plot. The data types on the horizontal and vertical axes comprise the two omic data types used in each trimodal model. The trifecta configuration allows for a better performance in both targeted and untargeted drugs, across almost all trimodal data combinations. Untargeted drugs remain easier to predict than targeted drugs across all configurations and data types.

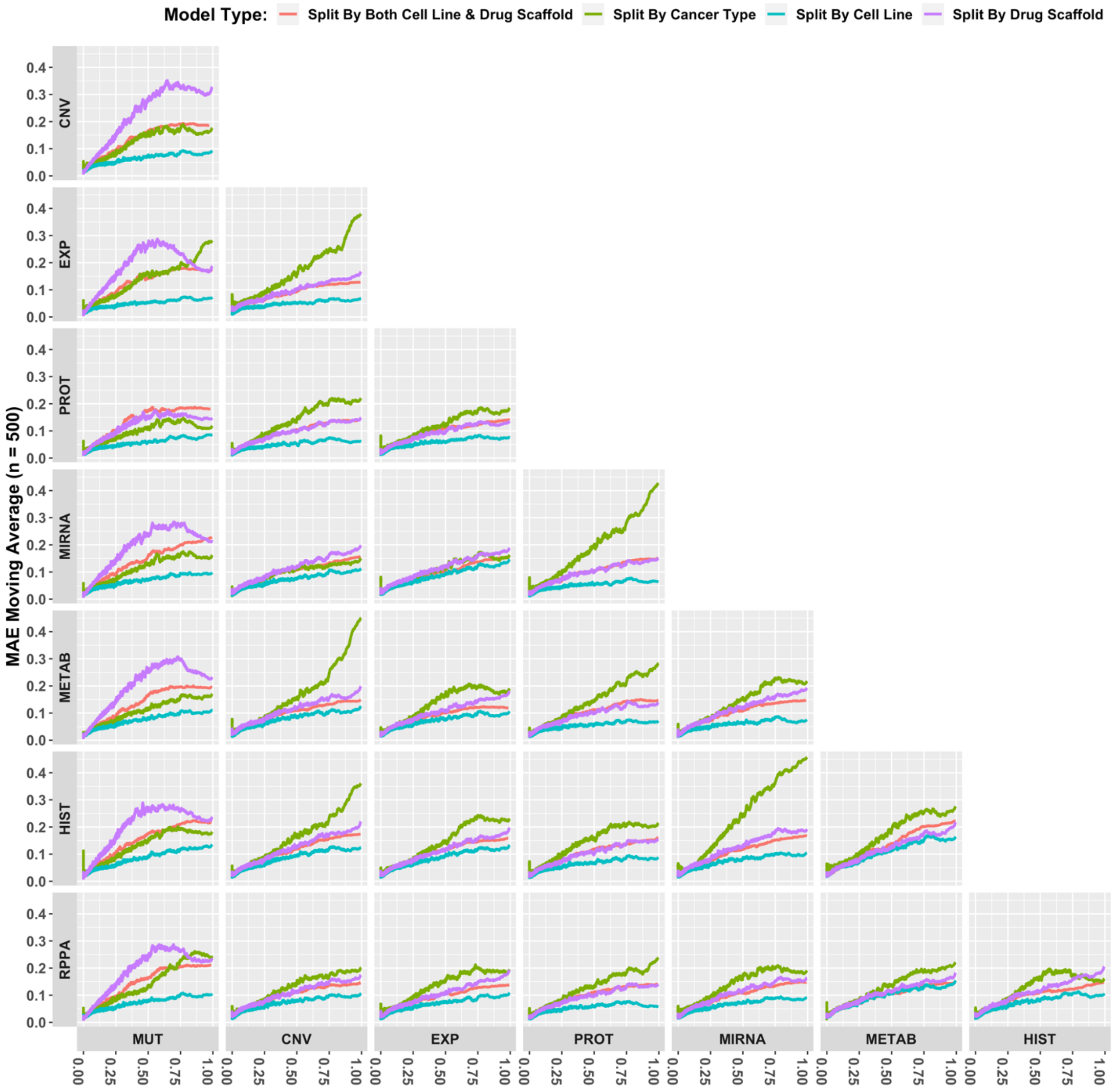


**Supplementary Figure 6. Comparison of different splitting methods in trimodal trifecta models.** Often, the Split by Cancer Type has the highest losses, indicating that the generalization to novel cancer types is the most difficult task among those tested.

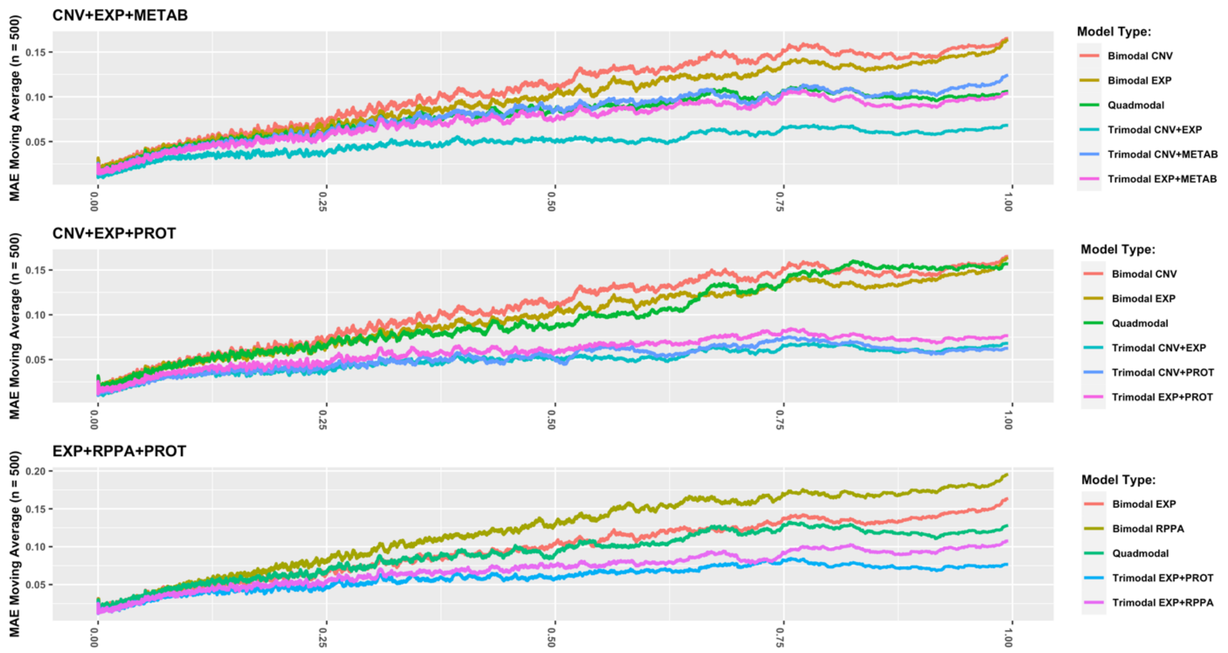


**Supplementary Figure 7. Comparison of quadmodal models with their trimodal and bimodal counterparts.** Trimodal models, which use two omic datatypes that comprise a subset of the three omic datatypes used by the quadmodal models, have a better performance.

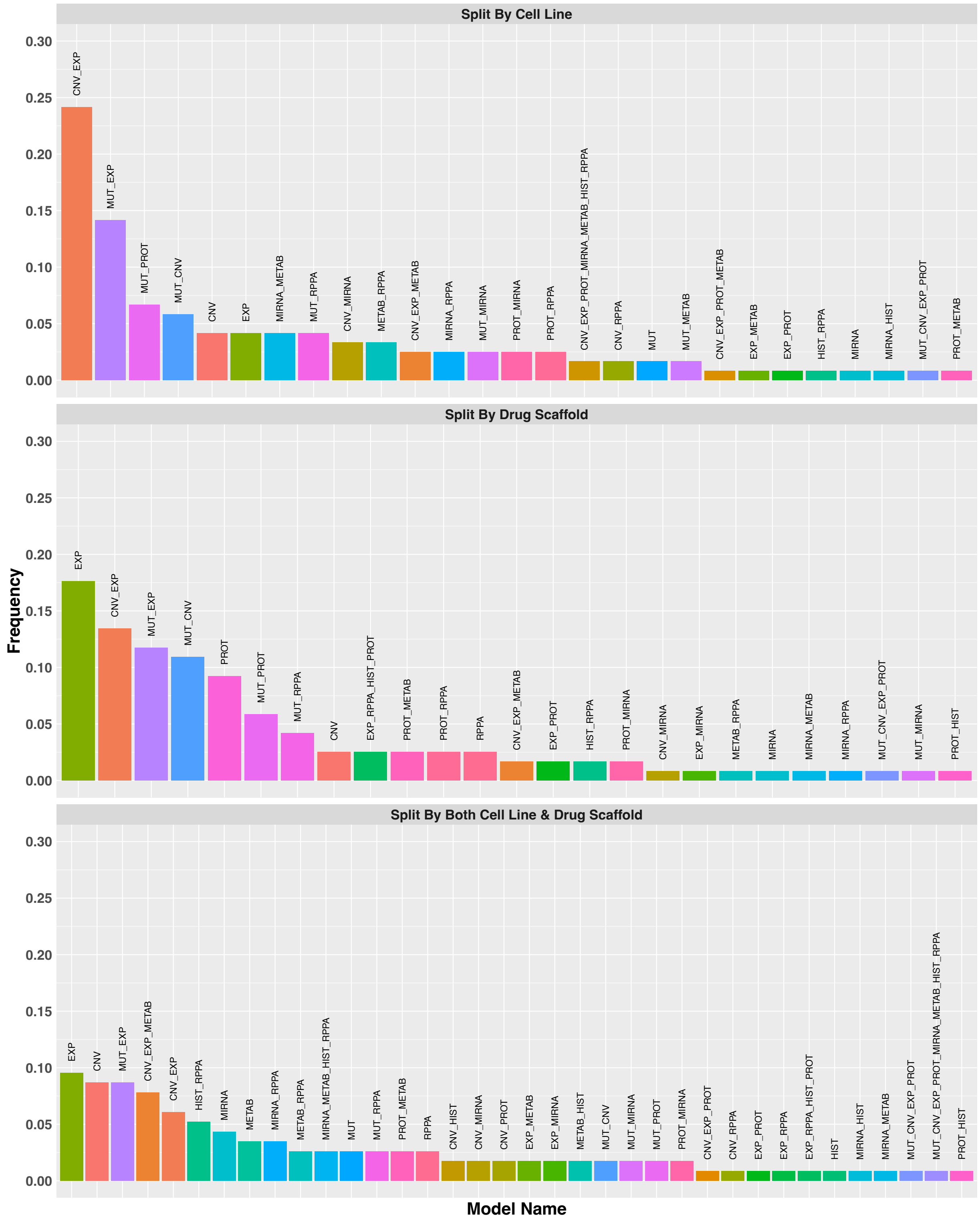


**Supplementary Figure 8. Frequency of omic data types used by models that predict AAC most accurately for each cell line and drug combination, divided by splitting schemes.**

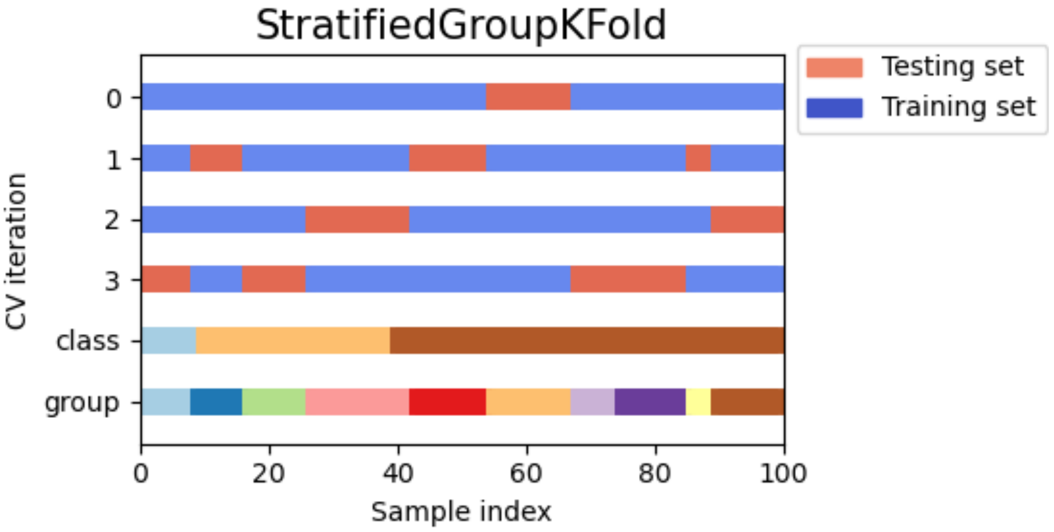


**Supplementary Figure 9. Stratified Group k-fold cross-validation data splitting scheme.**This scheme considers class and group distributions, and divides training and validation folds such that they more closely represent the original class and group distributions. This helps prevent biases in performance assessment. This figure was taken from: Visualizing cross-validation behavior in scikit-learn. https://scikit-learn.org/stable/auto_examples/model_selection/plot_cv_indices.html#sphx-glr-auto-examples-model-selection-plot-cv-indices-py[60]
