## Appendix for "Drug Response Prediction and Biomarker Discovery Using Multi-Modal Deep Learning"

### Data

#### Cell Line Characterization Data

Most of the cell line characterization data used in this work originated from the Cancer Cell Line Encyclopedia Project[1,2], which was merged into the Broad Cancer Dependency Map Project (DepMap) in 2018. DepMap version 21Q2 hosts all the biological data used in this project and can be openly accessed from https://depmap.org/portal/. Each data type is represented as a matrix, where the rows are cell lines, and columns are numerical values representing quantities of corresponding features. Because of the data incompleteness problem, each cell line profiling dataset has a different number of rows and columns. We used the cell line characterization data from DepMap, which has a large overlap with the cell lines used in the drug response dataset.

##### Point Mutation and Small Insertion-Deletion Data

Cell line mutational data in DepMap was initially assessed using massively parallel sequencing, focusing on 1,650 cancer-related genes[1]. Subsequently, Illumina sequencing-based whole-exome and whole-genome sequencing expanded mutational profiling to the entire genome[2]. Database maintainers have filtered results for germline variants. Single, Double, and Triple Nucleotide Polymorphisms (SNP, DNP, and TNP, respectively), as well as insertions and deletions, have been identified, resulting in 1.27 million mutations overall in 1,750 cell lines. These mutations overlap with mutations frequently observed in TCGA and COSMIC datasets, which are referred to as hotspots. To focus the analysis on conserved and potentially damaging mutations in addition to reducing the input features of the model, using DepMap's annotations, we selected genes that had mutations overlapping with TCGA and COSMIC hotspots, resulting in 8,752 and 91 genes, respectively, for a shared total of 8,772 genes. As a result, each cell line has an associated binary vector representing whether each gene has a hotspot mutation (1) or not (0).

##### Copy Number Aberration Data

Like point mutation data, CCLE copy number data was initially acquired using SNP microarrays and later complemented with whole-exome sequencing data. In summary, the genome is divided into segments, and a copy number is assigned to each segment by an algorithm based on the number of reads mapped to that segment. Moreover, gene-level copy numbers can be measured by mapping genes onto these segments. Again, DepMap has harmonized and preprocessed this data with log2 transformation with a pseudo-count of 1, providing copy-number profiles of 1,742 cell lines across 27,563 genes.

##### Gene Expression Data

CCLE gene expression data was initially assessed using GeneChip microarray technology[1] and subsequently using RNA-seq. This data provides transcript counts for protein-coding genes only and collapses the expression of all transcripts from the same gene (isoforms) into a single expression value. Reducing transcript-level expression to gene-level expression may reduce the amount of information captured from the cell[3], as different isoforms can result in different proteins with distinct functions. However, this also reduces the computational burden on statistical models. Gene expression data is preprocessed and harmonized by DepMap, resulting in Transcripts-per-Million (TPM) log2 transformed with a pseudo-count of 1, providing gene-expression profiles of 1,379 cell lines with 19,178 protein-coding genes.

##### Protein Quantification Data

High-throughput protein-quantification data has only recently become available for some cancer cell lines in the CCLE through work done by the Gygi lab[4]. The cell lines in this dataset are dominated by those from solid organs. Before applying mass-spectrometry to large-scale protein quantification, protein quantification was measured with reverse-phase protein arrays (RPPA), which are limited by throughput, coverage, and the availability of antibodies for proteins of interest. The Gygi lab used tandem mass spectrometry with three stages of separation ($MS^3$) with a quadrupole ion trap and tandem mass tag (TMT) labeling that allowed multiplexing of 10 samples. They also used a special normalization procedure described in a separate paper[5]. A bridge sample that contains an equal amount of each cell line used in the run is used for normalization. The normalized value for each protein is roughly the $\log_2$ ratio compared to the bridge; therefore, the values can be negative. Secondly, because the quantification uses relative values, each protein quantification value is only comparable with values for the same protein across samples and not to other quantitative values for other proteins. Although there are around 12,000 proteins quantified overall, only around 5,000 of them are observed across all cell lines with high confidence, with the rest missing in different cell lines, primarily due to lower abundance. While the difference among undetected proteins across different cell lines may be indicative of an underlying biological process, applying the remaining ~7,000 proteins as input to a machine learning model is a non-trivial problem. Therefore, we only used the subset of ~5,000 proteins that were detected in all cell lines in this project.

**MicroRNA (miRNA) Data**

MicroRNAs (miRNAs) are small RNAs (~22 nucleotides long) that post-transcriptionally regulate the expression of thousands of genes in a broad range of organisms, first discovered in 1993[6]. miRNA profiling data could be a better proxy for protein expression when combined with transcriptional data. Ghandi et al. used the hybridization-based NanoString platform to quantify the expression of 734 miRNAs across 953 cell lines[2].

**Metabolomics Data**

In cancer, the dysregulated activity of pathways can result in changes in cell metabolism. These changes then alter the quantities of metabolites that may be essential to normal cellular function. Therefore, characterization of the metabolome can help us understand cancer mechanisms better. Furthermore, some cancers have specific metabolic dependencies that have been directly targeted by therapies[7]. Using Liquid Chromatography-Mass Spectrometry (LC-MS), Li and Ning et al. partially characterized the metabolomes of 928 cell lines by quantifying 225 metabolites[8].

**Histone Modification Profiling Data**

The post-translational modification of histone proteins affects their function in nucleosomes that can ultimately remodel chromatin architecture, affecting the transcription of genes. Aberrant histone modifications have been associated with numerous diseases, including cancer. Thus, histone modification profiling data can better characterize cancer cells. Ghandi et al. used a mass spectrometry-based method to profile relative changes in the levels of almost all common post-translational modifications, including methylation and acetylation, phosphorylation, and ubiquityl modifications[2]. This resulted in 42 single and combined histone modifications across 896 cell lines.

##### Reverse Phase Protein Array Data

RPPA is essentially a dot-plot (similar to a western-blot) platform that allows the measurement of protein expression levels for multiple samples and proteins in parallel. Unlike mass-spectrometry-based methods, this method is dependent on high-quality antibodies for the proteins. Since not all proteins have suitable antibodies, only 214 proteins across 898 cell lines are quantified using this method.

#### Drug Data

Although representing mutations, copy numbers, or gene product quantities as numbers is intuitive, representing molecular compounds is more challenging. One widely used method to represent molecular compounds as numbers that computers can understand is by using molecular fingerprints. Fingerprinting can extract a molecule's structural properties and other features and represent them as a vector. It should be noted that vector similarity does not necessarily mean molecular similarity, as it is not yet canonically defined. Molecules can also be represented as SMILES which is one way of describing the structure of a chemical with ASCII strings, i.e., as a linear notation. The CTRPv2 dataset contains the SMILES representations of the drugs, which can be converted to a numerical molecular representation using the RDKit[9] package for cheminformatics. This molecular representation can then be used to calculate the extended-connectivity fingerprint or the molecular graph using the Pytorch Geometric[10] Python package. RDKit allows features such as aromaticity and hybridization for each molecule to be inferred from the SMILES representation.

In addition to representation, we have labeled the 481 drugs in CTRPv2 as targeted or untargeted to distinguish them in the performance assessment stage only. Targeted here refers to the subset of drugs that were rationally designed to inhibit specific genes and gene products involved in cancer-related pathways. The list of FDA-approved targeted drugs was obtained from https://www.cancer.gov/about-cancer/treatment/types/targeted-therapies/targeted-therapies-fact-sheet#what-targeted-therapies-have-been-approved-for-specific-types-of-cancer. 31 of these drugs match those in CTRPv2 and are summarized in **Supplementary Table 1**. Untargeted drugs in this context refer to chemotherapeutics with broad effects and drugs with an unknown target(s), comprising the remainder 450 drugs in the CTRPv2 dataset. Note that the model is not aware of whether a drug is targeted or not.

#### Drug Response Data

As discussed previously, the response of cells to drugs is often either measured by metabolic activity or DNA content. Both datasets used in this project use DNA content to measure the change in cellular viability. In addition, many cell lines are often only partially responsive to drugs with the concentrations tested; thus, the half-maximal inhibitory concentration ($IC_{50}$) is not experimentally observed and only algorithmically calculated. This issue compounds the problem of $IC_{50}$ being less descriptive than the AUC, as discussed in the introduction. Furthermore, when the dose-response curve is used in measuring the inhibitory potential of a drug, the lower the AUC is, the better. To make the interpretation of the response summary more intuitive, we use the area above the curve (AAC) instead.

We used the PharmacoGx package[11] to download and calculate AAC scores for the dataset from the Cancer Therapeutics Response Portal version 2 [12–14] (CTRPv2). It must be emphasized that this area above the curve (and the area under the curve) in this project only refer to the dose-response curve and are not to be confused with areas under and above the receiver operating characteristic (ROC) curve, which are used to show the performance of classification models.

The CTRPv2 project has aimed to reduce non-drug effects on cell growth by keeping cell media and other conditions consistent for all tested cell lines. This was done by maintaining nutrient levels and types, anti-microbial agents, temperatures, and oxygen levels. Furthermore, cell lines are assessed for cross-contamination or duplication by SNP-profiling panels. A summary of the data resources used in this project is provided in **Supplementary Table 2**.

Out of 310,792 samples in CTRPv2, 20,842 involve targeted drugs, and 289,950 involve untargeted drugs. **Supplementary Figure 3** compares the distributions of AACs from samples with targeted and untargeted drugs. A bigger proportion of samples with untargeted drugs have AACs closer to zero. Targeted drugs also have a slightly higher median AAC, 0.127 compared to 0.088 ($p$-value $< 2.22 \times 10^{-16}$, Mann-Whitney U Test).

### Methods

#### Standardization

This is also known as standardization, or Z-score normalization, and its benefits have been proven theoretically[15]. The training subset of each omic dataset is standardized column-wise (i.e., by each feature) by subtracting the mean (μ) and dividing by the variance (σ) in that subset **(Equation ( 1 ))**. The mean and variance are known as the Gaussian (normal) statistics and are used to scale the other remaining data points from each dataset accordingly. The gaussian statistics during cross-validation change as the training and validation subsets change.

$$Z = \frac{x - \mu}{\sigma} \qquad (1)$$

#### Label Distribution Smoothing

Ensuring that the model makes informed predictions rather than guessing the mean is crucial. To partially address the dataset imbalance, we have employed a sample weighting scheme inspired by classification problems. Intuitively, data points from less prevalent classes are weighted higher than those from more prevalent classes. Since drug response prediction is a regression task that does not contain classes, we divide the AAC distribution, which is in the [0,1] range, into bins of width 0.01. The difference with classification is that these bins, unlike classes, are more or less similar based on how close or far they are from each other, respectively. Using this knowledge, the label distribution smoothing (LDS) technique uses a symmetric kernel such as a gaussian kernel to smoothen the distribution **(Equation ( 2 ))**, making the class prevalence-based weighting scheme applicable to data used for regression, i.e., continuous data.

$$\boldsymbol{\tilde{p}}(\boldsymbol{\acute{y}}) \triangleq \int_{\boldsymbol{y}} \boldsymbol{k}(\boldsymbol{y}, \boldsymbol{\acute{y}})\boldsymbol{p}(\boldsymbol{y})\boldsymbol{dy} \qquad (2)$$

Where $\mathbf{k}(\mathbf{y}, \boldsymbol{\acute{y}})$ is a symmetrical kernel that satisfies the properties: $\mathbf{k}(\boldsymbol{y}, \boldsymbol{\acute{y}}) = \mathbf{k}(\boldsymbol{\acute{y}}, \boldsymbol{y})$ and $\nabla_{\mathbf{y}}\mathbf{k}(\mathbf{y}, \boldsymbol{\acute{y}}) = \nabla_{\acute{y}}\mathbf{k}(\boldsymbol{\acute{y}}, \mathbf{y})$ and $\mathbf{p}(\boldsymbol{y})$ is the number of appearances of label $\mathbf{y}$ in the training data, and $\boldsymbol{\tilde{p}}(\boldsymbol{\acute{y}})$ is the effective density of label $\boldsymbol{\acute{y}}$. Effectively, each bin now has an assigned factor corresponding to how populated it is. The inverse of this factor is then scaled and multiplied by the loss value during training **(Equation ( 3 ))**.

$$\boldsymbol{w_{\acute{y}}} = \frac{\mathbf{1}}{\widetilde{\boldsymbol{p}}(\acute{\boldsymbol{y}})}\ ,\ \mathbf{s} = \frac{\mathbf{100}}{\sum_{N=0}^{99} \boldsymbol{w_{\acute{y}}^{N}}} \qquad (3)$$

Where $\boldsymbol{w_{\acute{y}}}$ is the weight for a particular bin, and s is the scaling parameter that is multiplied by each weight. This process forces the model to emphasize learning from a specific data point based on how populated its bin and the neighborhood of bins are. This partially addresses the dataset imbalance problem and should make the model generalize better to data farther from the average. Note that this method is only used during training, and sample weights are not used during inference.

#### Graph Neural Networks

In this project, drug molecules are represented as graphs, where atoms are represented as nodes and bonds as edges. Nodes and edges can have properties such as aromaticity, hybridization, bond type, and chirality. GNNs then pass information from each node and its neighbors via its edges iteratively using smaller neural networks to embed either the atoms, the molecule, or both in the latent space. This embedding can then be used in an ANN for learning a specific task. In this project, We use the AttentiveFP GNN model[16] to extract features from drug molecules for the DRP task. As mentioned previously, We use the Pytorch Geometric[10] Python package to create the graph representations of drugs, which also implements the AttentiveFP model. One of the advantages of the AttentiveFP model is its use of the graph attention mechanism[17], which weighs neighboring nodes based on their relevance to the current node. The graph attention mechanism can be categorized into three operations of alignment, weighting, and context **(Equations ( 4 ), ( 5 ), and ( 6 ))**:

Alignment:

$$\boldsymbol{e_{vu}} = \boldsymbol{leaky\,relu}(\boldsymbol{W}.[\boldsymbol{h_v}, \boldsymbol{h_u}]) \qquad (4)$$

Weighting:

$$\boldsymbol{a_{vu}} = \boldsymbol{softmax}(\boldsymbol{e_{vu}}) = \frac{\boldsymbol{exp}(\boldsymbol{e_{vu}})}{\sum_{u \in N(v)} \boldsymbol{exp}(\boldsymbol{e_{vu}})} \qquad (5)$$

Context:

$$\boldsymbol{C_v} = \boldsymbol{elu}\left(\sum_{u \in N(v)} \boldsymbol{a_{vu}}.\boldsymbol{W}.\boldsymbol{h_u}\right) \qquad (6)$$

Where $\boldsymbol{v}$ is the target node, $h_v$ is the state vector of node $\boldsymbol{v}$, and $h_u$ is the state vector for the neighbor node $\boldsymbol{u}$. ***leaky relu*** and ***elu*** are variations of the ReLU activation function, which allow for small amounts of negative inputs instead of the floor of zero used in ReLU.

The AttentiveFP model itself is then formulated into two operations of message passing and readout (Equations ( 7 ) and ( 8 )):

Messaging:
$$\boldsymbol{C_v^{k-1}} = \sum_{\boldsymbol{u \in N(v)}} \boldsymbol{M^{k-1}}\left(\boldsymbol{h_u^{k-1}}, \boldsymbol{h_v^{k-1}}\right) \tag{7}$$

Readout:
$$\boldsymbol{h_v^k} = \boldsymbol{GRU^{k-1}}\left(\boldsymbol{C_v^{k-1}}, \boldsymbol{h_v^{k-1}}\right) \tag{8}$$

Where $\boldsymbol{h_v^k}$ is the state vector of target node $\boldsymbol{v}$ after $\boldsymbol{k}$ iterations and $\boldsymbol{N(v)}$ represents all the neighbors of node $\boldsymbol{v}$. The GRU function stands for the Gated Recurrent Unit[18]. The AttentiveFP model has outperformed other GNN architectures in most cheminformatic-related tasks. Therefore, we selected this architecture to extract contextual chemical information related to the DRP task.

In addition to the graph topology, nodes and edges have their own features fed to the GNN. Molecular features used in this project are summarized in **Supplementary Table 5** and **Supplementary Table 6**.

#### Low-rank Multimodal Fusion

As discussed before, to expand biomarker discovery to different facets of molecular data, devising a model that can process multiple 'omics inputs while addressing limitations in data completeness is crucial. Furthermore, different data types most likely bring complementarity rather than redundancy[19], especially in describing a group of cells at the molecular level, as observed when comparing quantitative proteomics data and transcriptional profiles[4]. The combination of multiple modalities can inform the development of novel clinical diagnostic tools.

Data integration in machine learning is often divided into early, intermediate, and late integration[20]. Early integration first combines various related datasets before they are used as input to a machine learning model. In contrast, late integration independently makes predictions using separate models for each data type, which are then used in an ensemble to make a final, aggregated prediction. The multi-modal learning approach used in this project follows an intermediate integration scheme, where each input data, whether biological profiling or drug-related data, is first embedded into a latent space by its respective encoder, and then the embedding, a.k.a. the code layer, is combined to create an integrated latent representation of all inputs as a single vector. This vector is then passed on to another neural network, which learns a mapping between these condensed representations and the drug response. Some approaches for combining the embeddings are discussed below.

Modeling interactions among the modalities and their embedded features reduces the ambiguities present when using unimodal models in sentiment analysis tasks. It can empower the model to make better predictions in the DRP task by drawing connections between different omic representations of the same cell line and drug embeddings. This is not possible using traditional data integration methods, namely integration by concatenation. One way to model these interactions is via a multi-linear pooling of each embedding. Bi-linear pooling is simply the outer product of two vectors (***M*** x 1 and ***N*** x 1), which results in an ***M*** by ***N*** matrix. The following shows the outer product of two 3 x 1 vectors ***u*** and ***v*,** which results in the 3 x 3 matrix ***A***.

$$\boldsymbol{u} = \begin{bmatrix} \boldsymbol{u_1} \\ \boldsymbol{u_2} \\ \boldsymbol{u_3} \end{bmatrix}, \boldsymbol{v} = \begin{bmatrix} \boldsymbol{v_1} \\ \boldsymbol{v_2} \\ \boldsymbol{v_3} \end{bmatrix}$$

$$\boldsymbol{u} \otimes \boldsymbol{v} = \boldsymbol{u}\boldsymbol{v}^T = \boldsymbol{A} = \begin{bmatrix} \boldsymbol{u_1 v_1} & \boldsymbol{u_1 v_2} & \boldsymbol{u_1 v_2} \\ \boldsymbol{u_2 v_1} & \boldsymbol{u_2 v_2} & \boldsymbol{u_2 v_3} \\ \boldsymbol{u_3 v_1} & \boldsymbol{u_3 v_2} & \boldsymbol{u_3 v_3} \end{bmatrix}$$

This is in contrast with the integration by concatenation, where multiple vectors are joined to create a single vector of elements:

$$\boldsymbol{u} = \begin{bmatrix} \boldsymbol{u_1} \\ \boldsymbol{u_2} \\ \vdots \\ \boldsymbol{u_N} \end{bmatrix}, \boldsymbol{v} = \begin{bmatrix} \boldsymbol{v_1} \\ \boldsymbol{v_2} \\ \vdots \\ \boldsymbol{v_M} \end{bmatrix}, \boldsymbol{s} = \begin{bmatrix} \boldsymbol{s_1} \\ \boldsymbol{s_2} \\ \vdots \\ \boldsymbol{s_D} \end{bmatrix}$$

$$\boldsymbol{J} = \begin{bmatrix} \boldsymbol{u_1} \\ \vdots \\ \boldsymbol{u_N} \\ \boldsymbol{v_1} \\ \vdots \\ \boldsymbol{v_M} \\ \boldsymbol{s_1} \\ \vdots \\ \boldsymbol{s_D} \end{bmatrix}$$

Multi-linear pooling extends this concept to more than two vectors, resulting in a tensor, a higher-dimensional form of a 2-dimensional matrix. The tensor fusion method also models the subset of the combinations of each modality (e.g., with four modalities, unimodal, bimodal, and trimodal sub-tensors are also modeled). A variant of this approach reduces the computational burden by using a low-rank approximation of the fusion tensor, which also can have modality-specific factors. This way, low-rank tensor fusion models interactions and can assign context-specific weights for each modality, addressing problems with concatenation and weighted sum methods.

### Hyper-parameter optimization

In this work, We use the HyperOpt[21] algorithm from the Ray[22] package's Tune[23] library, which can run trials in parallel, significantly speeding up the tuning process. While searching for optimal hyperparameters is a computationally intensive process, it ensures the most effective use of available data.

"Training hyper-parameters" refers to those parameters that control the speed of learning. We tuned both the learning rate and the batch size. Architectural hyper-parameters often outnumber training hyper-parameters, and in this project included: 1) the number of layers, 2) the number of neurons at each layer, 3) the activation function used at all layers, 4) whether batch normalization and dropout are used. For the last (output) layer, the sigmoid activation function without batch normalization or dropout is always used to ensure that the predicted AAC has a value between 0 and 1. Using the same activation functions, batch normalization, and dropout parameters for the whole model instead of layer-by-layer hyperparameters simplifies and shortens the tuning process as fewer hyperparameter combinations have to be explored. However, this grouping of hyperparameters may lead to non-optimal results, as it is essentially restricting the model architecture. To save time, we grouped the hyperparameters and relied on empirical literature results to manually select a few of these hyperparameters, such as the Swish activation function, which was discussed above.

Since omic autoencoder modules are first trained on all their respective available data, they also undergo the hyperparameter optimization process individually first. Once suitable hyperparameters for each module are found, the encoder subnetworks are extracted and attached alongside the drug encoder to the DRP module. Then, another hyperparameter search is performed only for the DRP module and the tensor fusion mechanism. The already optimized encoders are further trained under the DRP context. This is represented visually in **Figure 2**.

Each hyperparameter optimization trial uses grouped and stratified cross-validation for the whole model (i.e., the autoencoders and the DRP module). The average validation losses from the K folds are used as a final score for each trial. This can result in minor information leakage between the cross-validation and hyperparameter optimization processes. The hyperparameter optimization can aim to optimize the average cross-validation loss but is not then tested on an independent testing set. This can make the averaged cross-validation loss slightly less representative of real-world generalization performance, resulting in the underestimation of the losses. This happens mainly when many trials are run. A nested cross-validation scheme can alleviate this but is significantly more computationally demanding and time-consuming. We skipped the use of this method as We limited the number of trials to 40 for each model type.

#### Stratified Group K-fold Cross-validation

To test the predictive performance of my model on unobserved data, we used k-fold cross-validation[24]. This method divides the dataset into **k** bins, using each bin once as a validation set while using the other bins for fitting an untrained and identically initialized copy of the model. This results in training and validating the model independently **k** times, giving me a more accurate performance assessment of the generalization capabilities of my model compared to using only a single testing set. This method is non-exhaustive, meaning that not all possible subsets of data are used as training or validation. Since the distribution of cell lines by cancer types in DepMap is not uniform, training and validation sets might have unequal distribution of different cancer types, making evaluation less representative of real-world performance. To ensure the similarity of the distribution of data points between training and validation sets and to assess my models' generalization, we used stratification and strict subsetting, respectively. Stratification is a process to ensure that each bin or fold has the same proportion of cell lines based on their lineage as the entire dataset. This ensures that the model is trained and validated on data not biased towards a particular lineage or set of cell lines. Strict subsetting (a.k.a. grouping) differs from lenient subsetting in that it ensures that either the cell lines, drugs, or both are not used in both the validation and the training set, ensuring that the model will be validated on drug response data from cell lines or drugs that it has never seen before. The combination of stratification and grouping better assess the model's generalization performance to novel scenarios. We use the implementation of Stratified Group K-fold from the scikit-learn python package[25] (**Supplementary Figure 9**).

#### Interpretation using Integrated Gradients

There are different classes of ML interpretation techniques based on various criteria. Some models, such as shallow decision trees and sparse linear models, are intrinsically interpretable by analyzing internal model parameters. Other models require post-hoc interpretation, which is done by analyzing the model after training[26]. Furthermore, interpretation can be model-specific or model-agnostic, and some methods also seek to explain a complex model through a simpler, more interpretable conjugate model. A few well-known interpretation methods are provided in the Captum package for Pytorch[27], which we have used. Specifically, For this task, we used the Integrated Gradients technique[28], which is more suitable for neural networks. We then applied this interpretation method to identify important biological features and drug characteristics.
